# Data-driven analysis of motor activity implicates 5-HT2A neurons in backward locomotion of larval *Drosophila*

**DOI:** 10.1101/257568

**Authors:** Jeonghyuk Park, Shu Kondo, Hiromu Tanimoto, Hiroshi Kohsaka, Akinao Nose

## Abstract

Rhythmic animal behaviors are regulated in part by neural circuits called the central pattern generators (CPGs). Classifying neural population activities correlated with body movements and identifying the associated component neurons are critical steps in understanding CPGs. Previous methods that classify neural dynamics obtained by dimension reduction algorithms often require manual optimization which could be laborious and preparation-specific. Here, we present a simpler and more flexible method that is based on the pre-trained convolutional neural network model VGG-16 and unsupervised learning, and successfully classifies the fictive motor patterns in *Drosophila* larvae under various imaging conditions. We also used voxel-wise correlation mapping to identify neurons associated with motor patterns. By applying these methods to neurons targeted by *5-HT2A-GAL4*, which we generated by the CRISPR/Cas9-system, we identified two classes of interneurons, termed Seta and Leta, which are specifically active during backward but not forward fictive locomotion. Optogenetic activation of Seta and Leta neurons increased backward locomotion. Conversely, thermogenetic inhibition of *5-HT2A-GAL4* neurons or application of a 5-HT2 antagonist decreased backward locomotion induced by noxious light stimuli. This study establishes an accelerated pipeline for activity profiling and cell identification in larval *Drosophila* and implicates the serotonergic system in the modulation of backward locomotion.

## Introduction

The neural circuits generating rhythmic behaviors such as walking and breathing are called the central pattern generators (CPGs)^1–3^. Since rhythmic behaviors of invertebrates and vertebrates share many features, studies on the CPGs in one animal species are expected to serve as a model for those in other species^4^. Identification of neurons involved in CPGs is the first important step in understanding how the rhythmic behavior is generated and regulated by the neural circuits. Such analyses have often been performed in the isolated central nervous system (CNS) since it is known that CPGs can produce fictive motor patterns that resemble the actual behavior patterns without any sensory feedback^5–7^.

Recent advances in imaging technology such as spinning-disc and light-sheet microscopy enabled recording of neural activity in large regions in the brain, paving new ways to investigating CPG circuits. In animals with relatively small CNS such as *C.elegans*, larval *Drosophila* and larval zebrafish, it is now possible to image the entire brain or even the whole animal in real time^8–10^. While the technological advances are now enabling one to record the activity of a majority of neurons in the nervous system in these animals, it remains challenging to extract useful information from the large data-sets obtained by the recording. In the case of CPG studies, for instance, one may want to determine the time windows in which specific motor activity takes place and then to identify the neurons that show activity related to the initiation, duration, and termination of the motor pattern. Previous studies used methods such as principal component analysis (PCA), independent component analysis (ICA), singular-value decomposition (SVD) and *k*-means clustering to quantitatively classify neural activity patterns^8, 11, 12^. Although these methods are powerful and have been used widely, they often require manual optimizations which are labor-intensive and confined to specific datasets. Therefore, development of alternative methods which can handle various types of activity data more efficiently is greatly anticipated.

The larval *Drosophila* is one of the most powerful model systems for studying neural circuits related to rhythmic behaviors since its CNS is numerically simple (containing ∼ 10,000 neurons) and amenable to various genetic manipulations. Especially, imaging fictive motor patterns in the isolated CNS with genetically encoded Ca2+ indicators is well established^13^. An isolated CNS can generate fictive motor outputs such as coordinated propagation of motor activity along the body axis, which resembles forward and backward locomotion of the animal, and left-right asymmetric bursts in anterior neuromeres which likely correspond to turning^14^. Whole-animal functional imaging in embryos just before hatching^9^ confirmed that the propagating activity and asymmetric bursts occur during forward/backward locomotion and turning, respectively. An isolated CNS also generates symmetric and synchronous bursting activity in the anterior-most and posterior-most segments, which often but not always occur just prior to the initiation of backward and forward fictive locomotion, respectively^13^. While corresponding larval motor outputs of the bursting activity is not clear, bursts in posterior-most segments may be related to movement of the gut and tail which is known to occur prior to forward locomotion^15^. Previous studies have shown that subsets of interneurons show activity correlated with these fictive motor outputs and play roles in the regulation of larval movements, such as segmental activity propagation, left-right symmetric coordination and differential recruitment of motor pools^16–22^.

In this study, we present a new methodology for classifying neural activity patterns in larval *Drosophila* that utilizes a convolutional neural network (CNN) and unsupervised learning. This method successfully classified forward and backward waves as well as synchronous activities in anterior- and posterior-most neuromeres from large activity data derived from different sub-populations of central neurons. We also identified cells associated with the classified motor activity patterns by voxel-wise correlation mapping. We then applied this method to a population of neurons targeted by *5-HT2A-GAL4*, which we generated by CRISPR/Cas9-mediated gene knock-in^23^, and successfully identified two classes of interneurons that are specifically active during fictive backward locomotion. Further functional analyses implicated these *5-HT2A* neurons and 5-HT neurons in the regulation of backward locomotion. Our study provides an accelerated pipeline that efficiently identifies neurons correlated with motor patterns from wide-volumetric functional imaging data.

## Results

### Classification of fictive motor patterns by deep convolutional feature and unsupervised clustering

In this study, we aimed to identify motor patterns automatically in a data-driven manner. The key idea was to create x-t images in which x-axis represents the activity of multiple ROIs at each time point. Since the motor activity pattern in the x-t image is similar to images such as handwritten digits^24^, we expected that CNN, known to give superior performance in image classification tasks^24–26^, may be used to efficiently categorize the motor patterns. We, therefore, tested if a CNN model, VGG-16 pre-trained on the ImageNet dataset^27, 28^, which is widely used in transfer learning^29^, can classify neural activities from the calcium imaging data. We used *RRa-GAL4* to express the genetically encoded calcium indicator GCaMP6f in aCC and RP2 motoneurons (MNs) in third instar larvae and imaged fluorescence changes associated with neuronal activity in the isolated CNS using spinning-disc confocal microscopy. As previously reported^13, 16^, the MNs showed left-right symmetric wave-like activity patterns reflecting forward and backward locomotion (hereafter called FW and BW, respectively) and bilateral asymmetric activities associated with turning. We used neural activity data from 18 regions of interest (ROIs) corresponding to both sides of the neuromeres T2-A7 in 12 animals for the analyses (Fig. 1a). Since our initial goal was to identify motor patterns corresponding to FW and BW, which are bilaterally symmetric, we initially combined the activity data in the left and right side and extracted activity patterns associated with propagating waves, and only later used the information from both sides to study the symmetry of the motor patterns identified. Hence, we compressed the 18-dimension (18-d) time series to 9-d time series using the maximum value of each segment (Fig. 1b, 2a) and analyzed the data using the CNN model (Fig. 1c, S1a).

**Figure 1.**
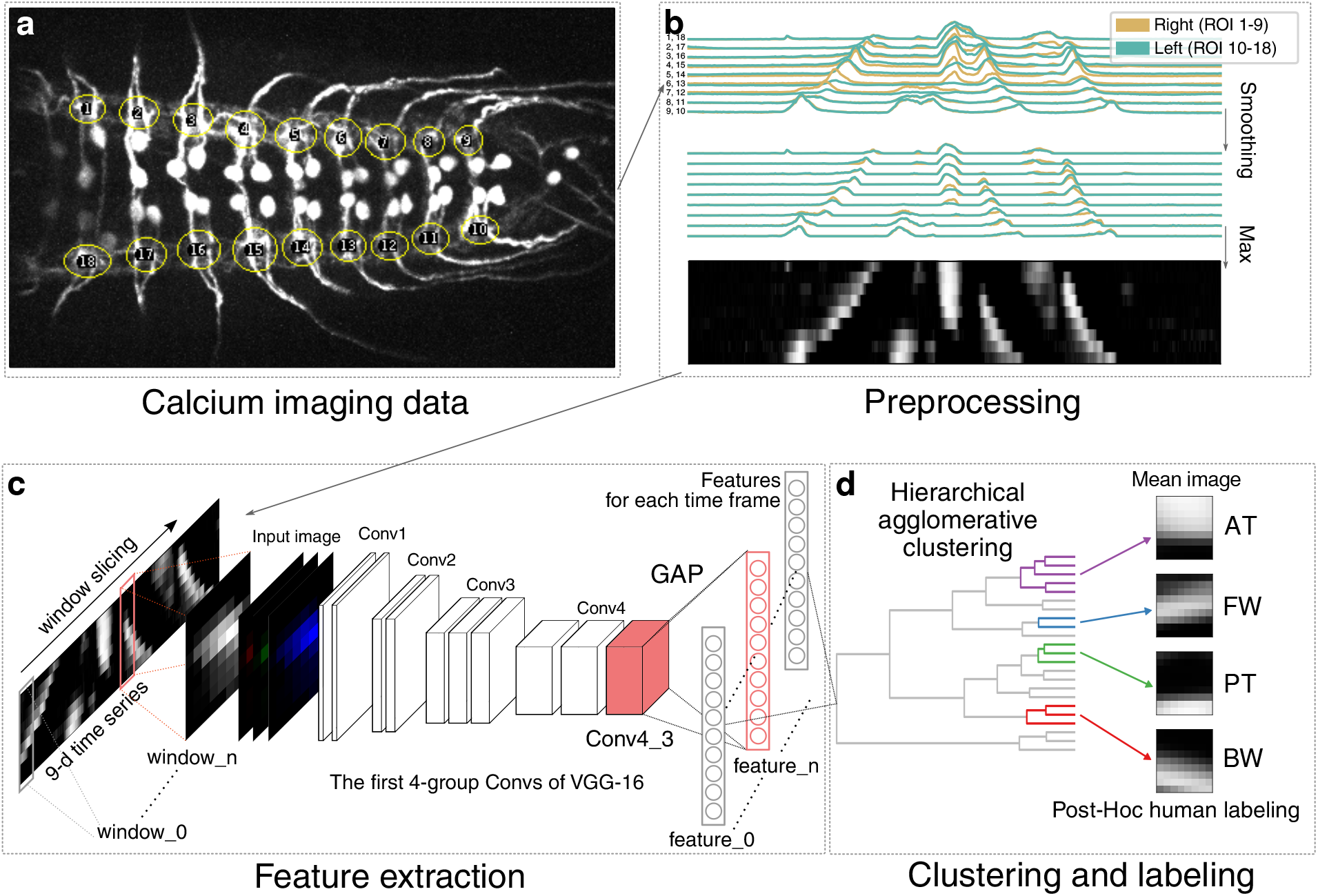
Outline for motor pattern classification using deep convolutional feature. (a) Neural activity of segmental aCC MNs was captured in 18 ROIs, covering each side of the neuromeres T2 - A7 (visual cues). (b) Preprocessing of the activity data. The time series representing the activity in each ROI was smoothed to reduce the effects of degeneration. Then, the data on the left and right side of each neuromere was compressed by adopting the maximum value to 9-d time series. (c, d) Framework for automated identification of motor patterns. (c) The 9-d time series was decomposed into windows with constant intervals. The windows were converted into a square 8-bit gray-scale images and then to RGB images. Features of each RGB image were extracted from the 3rd layer of the 4th convolutional block (Conv4 3) of VGG16 by global average pooling. (d) The features are classified into clusters by hierarchical agglomerative clustering (HAC) using Ward’s method. See Fig. S1 for details.

**Figure 2.**
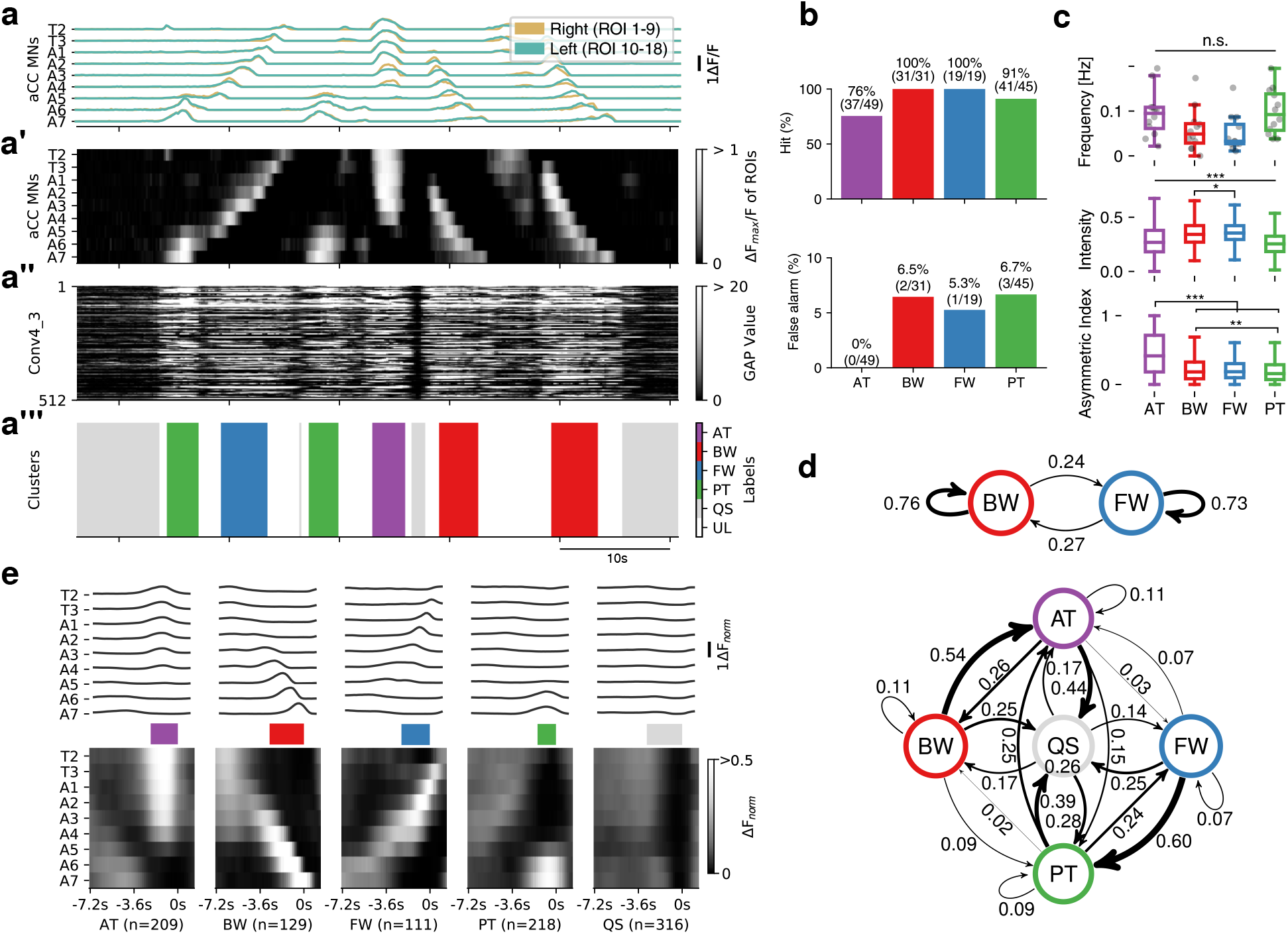
Evaluation and characterization of fictive motor patterns classified by deep convolutional feature. (a) An example of extracted motor patterns (a”’) from the activity data (a) Intermediate layers, the pre-processed 9-d time series displayed as a grayscale image (a’) and global average pooled values of Conv4 3 (a”), are also shown. AT, anterior burst; BW, backward wave; FW, forward wave; PT, posterior burst; QS, quiescence state; UL, unlabeled, represented in different colors as shown in the color-code bar. T2-T3, thoracic segment 2 to 3; A1-A7, abdominal segment 1 to 7. See Fig. S1a for details. (b) Evaluation of the extracted motor patterns. Hits (top) and false alarm (bottom) for the four motor patterns are shown. (c) Frequency (top), intensity (middle) and symmetry (bottom) of the four motor patterns extracted from 12 samples. Intensity represents the maximum fluorescence change (∆F*_norm_*) among all ROIs during each motor pattern. Asymmetric index is defined as the difference of the maximum ∆F*_norm_* in the left and right ROI divided by the maximum ∆F*_norm_* in all ROIs at each time-frame. \**p* 0.05, ** *p* 0.01, *** *p*0.001, ANOVA with post-hoc Tukey HSD test. (d) Markov chain of fictive motor patterns. Transition probabilities derived from 2 states (BW and FW, top) and 5 states (AT, BW, FW PT and QS, bottom). Thickness of arrows represents the transition probability between the motor patterns. Note that the arrow from QS to QS is omitted due to lack of space. (e) Averaged fictive motor patterns. Line plot (top) and grayscale image (bottom) are shown aligned with the end points of each motor pattern. Colored bars in the middle represent the average duration of each motor pattern (AT:2.2 ±0.1s, BW:2.8 ±0.1s, FW:2.3 ±0.1s, PT:1.5 ±0.1s, QS:2.9 ±0.3s, mean± standard deviation).

We searched for the layer which gives the best match for FW and BW by cluster analyses of extracted features from each layer of the CNN model with hierarchical agglomerative clustering (HAC, Ward’s method)(Fig. S1b, c),. By consulting the original activity pattern, we manually determined the clusters representing FW and BW, and found that the intermediate Conv4 3 layer gives the best performance in classifying the motor patterns from the neural activity data (Fig. 1c, d, Fig. S1). In addition to activities reflecting FW and BW, the Conv4 3 layer also succeeded in identifying two additional motor-related activities described previously^13^. One is left-right symmetric or asymmetric activation in anterior neuromeres (hereafter called AT). The other is synchronous and bilateral activation in posterior-most neuromeres (called PT) and is known to often occur before forward waves^13^. It was difficult to correlate these activities to motor patterns in intact larvae. While some of the AT activities likely correspond to turning, we were unable to clearly distinguish among turning-like and other AT activities. Furthermore, the classification performance decreased when asymmetric activities were taken into consideration (i.e., using 18 ROIs). From these reasons, we decided to focus on classifying activity patterns without considering asymmetric parameters in this study and took the maximum value in each segment so that we can detect activities occurring in either side of the animal. We evaluated the efficacy of the identification by comparing with the ground truth data obtained manually (∼10% of dataset) and found that our methodology yields on average, 75-100% correctly classified hits and 0-6.7% false alarms for each the four motor patterns, FW, BW, AT and PT (Fig. 2a-a”’, b, S1d). It is noteworthy that the CNN layer followed by HAC analyses succeeded in extracting all four major motor-related activities identified by human inspection (Fig. 1d)^13^. In the following analyses, we also annotated the time windows with no activity as a quiescent state (QS) and those showing activities other than the four motor patterns as unlabeled (UL).

We characterized the frequency and bilateral symmetry of the four motor patterns identified (Fig. 2c). The frequency of bursting activities in anterior and posterior neuromeres (AT and PT) and wave-like activity (FW and BW) were not statistically different. Intensities of neuronal activation were found to be higher during BW and FW compared to AT and PT. We also confirmed that AT but not other motor patterns shows strongly asymmetric activity as previously reported^13^. It has been known that transition between different motor patterns in behaving larvae is biased^30, 31^. We, therefore, asked whether the transition probabilities of fictive behavior patterns are also biased and found it is indeed the case (Fig. 2d). Strongest transition probabilities were seen in the transition from BW to AT and that from FW to PT (Fig 2d, bottom). Since the transition probabilities from AT to BW and from PT to FW (directly or via QS) were also high, the probability of the occurrence of the same motor pattern was high after BW and FW (i.e., BW-AT-BW, FW-PT-FW) (Fig 2d, top). While direct transition between AT and PT frequently occurred (0.15-0.25 transition probabilities), direct transition between BW and FW did not occur. These tendencies (e.g., AT occurs before BW) were also observed in the average image of each activity (Fig. 2e). This is consistent with the fact that larvae often perform forward or backward locomotion consecutively. Thus, our methodology successfully classifies motor patterns and allows efficient quantification of a large-scale activity data without laborious human labeling processes.

### Voxel-wise mapping of whole neurons

In the above experiment, GCaMP6f was expressed in a small number of identified MNs in each segment. We next asked whether the semi-automated classification method of fictive motor patterns can be applied to imaging data from a larger number of cells and, if so, whether we can use the classification method to identify previously uncharacterized interneurons that show activity pattern(s) correlated with specific fictive motor pattern(s). For this purpose, we first used a pan-neuronal GAL4 driver, *elav-GAL4*, to express GCaMP6f in all neurons. As with the above experiment, we analyzed the activity extracted from 18 ROIs representing all hemisegments in three larvae and succeeded in extracting the four motor patterns (Fig. 3a, b, S2a). Thus, our methodology is also applicable to pan-neuronal activities. We next tried to identify voxels correlated with each of the fictive motor patterns by using Pearson’s correlation coefficient (Pearsons’ r, Fig. 3c-c””). Since some voxels showed correlation with multiple motor patterns, we also determined the motor pattern which showed the highest correlation with the signal in each voxel (Dominant motor pattern map, Fig. S2b-d). Voxels correlated with AT (>0.1 Pearson’s r) were found in large quantities in anterior neuromeres (T1 to A3, Fig. 3c’), whereas those correlated with BW were seen in a complementary manner in posterior neuromeres (A4 to A7, Fig. 3c”). AT was often seen prior to BW and these two motor patterns together appear to correspond to the backward locomotion of the larvae. Similarly, voxels correlated with FW were largely found in middle neuromeres (A1 to A5, Fig. 3c”’), whereas those correlated with PT were seen in a complementary manner in posterior neuromeres (A6 to A8, Fig. 3c””). PT was often seen prior to FW and these two motor patterns together likely correspond to the backward locomotion of the larvae. These voxels were seen in similar manners across individuals (Fig. S2b-d) and likely correspond to segmentally repeated neurons showing propagating activity (FW and BW) or bursting activities spanning several segments (AT and PT). We also identified voxels that were separated from the segmentally clustered ones described above and showed unique morphologies and thus likely belong to the same cell(s). These included structures in a dorso-medial region in anterior neuromeres which showed correlation with BW (Fig. 3c”, arrow) and a pair of longitudinal axons in dorso-lateral neuropile, likely those of ascending/descending neurons, which showed correlated activity with PT (Fig. 3c””, arrow). By placing a ROI in each of these structures (Fig. 3c”, c””), we confirmed that they are active specifically in BW or PT (Fig. 3d)

**Figure 3.**
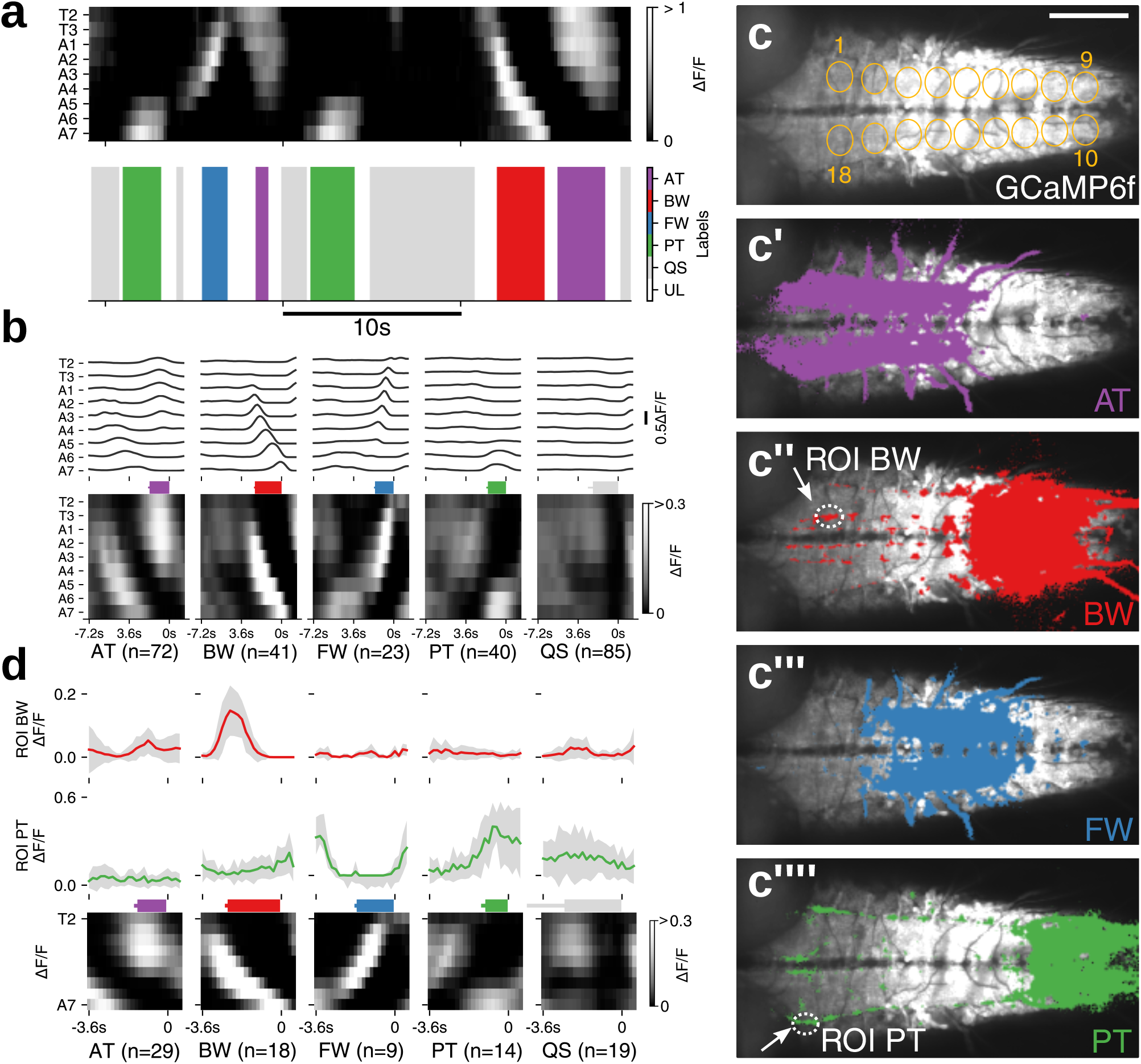
Motor pattern extraction from activity data of whole neurons. (a) An example of extracted motor patterns (bottom) from the activity data derived from pan-neuronally expressed GCaMP6f (top, grayscale image) (b) Mean of fictive motor pattern dervied from 3 samples aligned with the end point of each label. Average duration of each motor pattern, AT:1.8 ± 0.1s, BW:2.4 ±0.1s, FW:1.7 ±0.1s, PT:1.6 ±0.1s, QS:2.3 ±0.4s (mean±standard deviation). Note that unnormalized ∆F/F is shown since normalization obscured the motor pattern. (c) Motor pattern correlation mapping. Voxels with *>*0.1 Pearson’s r for each motor pattern, AT (c’), BW (c”), FW (c”’) and PT (c”’), are shown in colors. c shows the 18 ROIs used for the analyses (visual cues). Image baselines are derived from maximum GCaMP signal. (d) Neural activity in the structures identified by the correlation mapping. The average activity in the structures identified as specific to BW (top) and PT (middle) during each motor pattern (bottom). Scale bar 100 *µ*m.

### Voxel-wise mapping of putative 5-HT2A-receptor expressing cells identified Seta and Leta as backward-specific neurons

The correlation mapping of motor patterns described above enabled us to identify neuronal activities specifically associated with each of the four motor patterns. However, since all neurons expressed GCaMP6f, it was difficult to identify the cells that showed these activities. To overcome these limitations, we next tried to use GAL4 lines that drive expression in subsets of neurons. In particular, we were interested in neurons expressing neuromodulator (NM) receptors because of their potential roles in behavioral control. We therefore used the CRIPSR/Cas9 method to generate new GAL4 lines that recapitulate the expression pattern of each NM receptor. In these lines, the *2A-GAL4* gene^23^ was inserted in frame with each NM receptor immediately in front of the stop codon, allowing GAL4 to be co-expressed with the NM receptor in the same spatiotemporal pattern (Fig. 4a). We generated *GAL4* knock-in lines for 12 NM receptors (serotonin-receptors, *5-HT1A, 5-HT1B, 5-HT2A, 5-HT2B, 5-HT7*; octopamine-receptors, *Octβ 2R, Octβ 3R, OamB*; tyramine-receptor, *TyrR2*; dopamine-receptors, *Dop1R1, Dop2R, DopEcR*) and studied the expression of each GAL4 line in larvae using *UAS-mCD8-GFP*as a reporter. Except for *5-HT2A-GAL4*, all of the NM receptor GAL4 lines drove GFP expression in large subsets of neurons in the larval CNS (Fig. S3). In contrast, *5-HT2A-GAL4* drove GFP expression in small subsets of neurons which can be individually identified (Fig. 4b). We performed calcium imaging of the cells expressing *5-HT2A-GAL4*, analyzed the data in a data-driven manner as described above (as a reference for motor activity, we also expressed GCaMP6f in aCC and RP2 MNs) and generated a dominant motor pattern map (Fig. 4c, c’). By this, we identified voxels which were strongly active during BW but not FW locomotion (Fig. 4c”, S4d). As will be described below, we found these voxels belong to two pairs of interneurons in the VNC, named as Seta (Small-*η*) and Leta (Large-*η*) after their characteristic morphologies (see below). Although Seta and Leta neurons were both active during BW (Fig. 4d), temporal profiles of the activity of these neurons differed: while Seta showed a single peak of activity during BW, Leta showed double peaks. Furthermore, Seta but not Leta showed weak activity during AT and PT (Fig. 4e).

**Figure 4.**
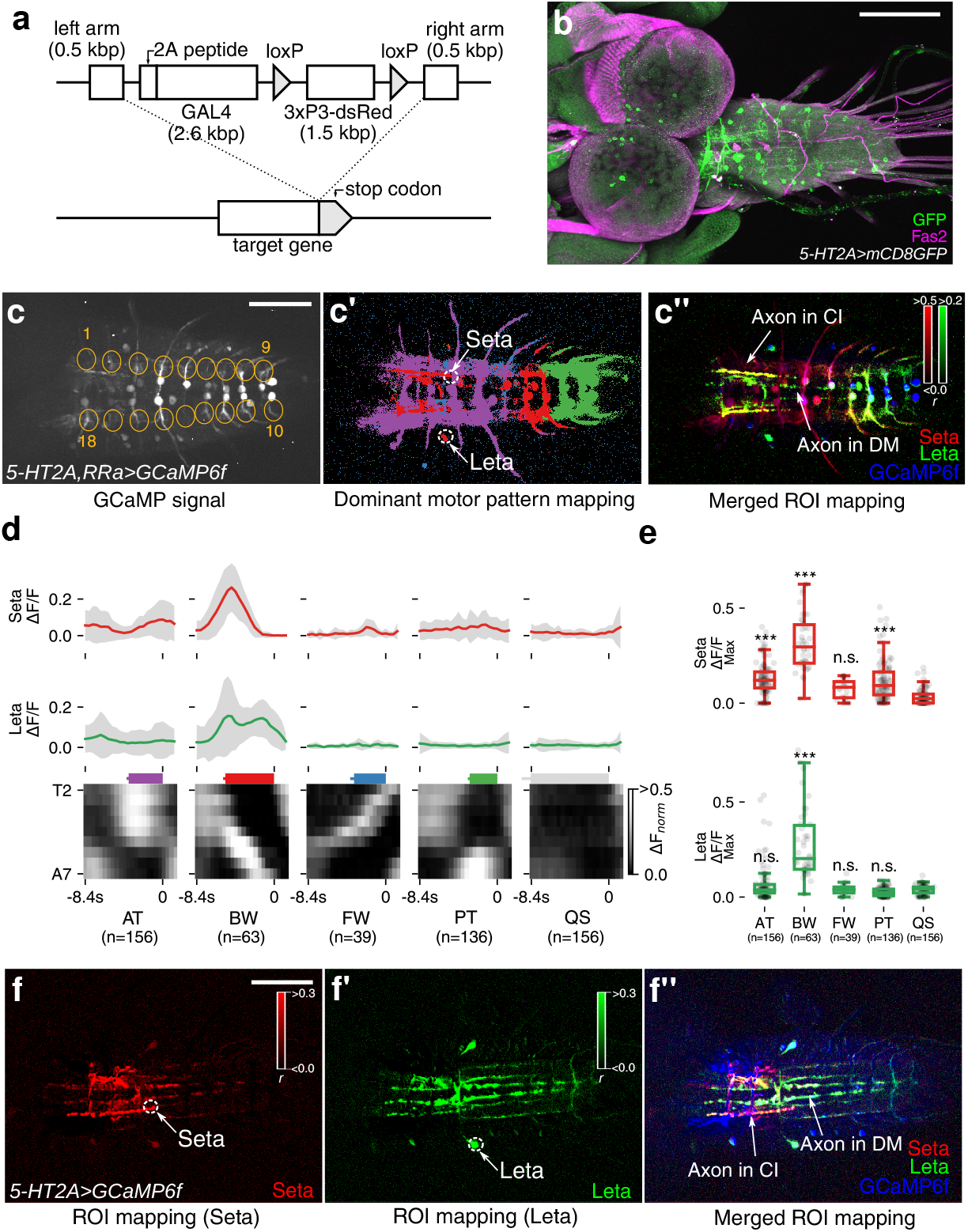
Identification of Seta and Leta, backward-specific interneurons expressing *5-HT2A-GAL4*. (a) A cartoon showing the design of GAL4 knock-in. The GAL4-coding sequence along with the 2A peptides was inserted by the CRISPR/Cas9 method to the target NM receptor gene locus, just downstream of the C-terminus of the NM coding sequence. dsRed flanked by loxP sites was used as a marker for successful insertion. (b) Expression of *5-HT2A-GAL4*visualized with *mCD8-GFP*. (c-c”) motor pattern mapping in *5-HT2A-GAL4*- and *RRa-GAL4*-targeted cells. (c) 18 ROIs representing each hemisegment used for the analyses (visual cues). (c’) Dominant motor pattern mapping identified voxels specific to BW. Motor patterns are colored as in Fig. 3. Two ROIs (labeled as Seta and Leta) are selected for the ROI correlation mapping in (c”). (c”) ROI correlation mapping reveals the outline of Seta and Leta. Pearson’s r between each voxel and a ROI in Seta and Leta shown in (c’) was calculated. The mapping results for Seta (red) and Leta (green) and GCaMP signals (blue) are merged. Axons of Seta and Leta are revealed in yellow (red+blue) and green/blue color (green+blue) and are found to be present in the CI and DM tracts (Fas2 coordinate33), respectively (arrows). c’ and c” are z-stacked images. (d) Mean activity of Seta (top, red) and Leta (middle, green) (derived of three samples) aligned with motor activities during AT, BW, FW and PT (bottom, grayscale image). Colored bars represent the duration of each motor pattern. (AT:3.6 ±0.2s, BW:5.2 ±0.2s, FW:3.4 ±0.4s, PT:3.0 ±0.1s, QS:8.4 ±0.9s, mean±standard deviation). (e) Maximum activity level (fluorescence changes, ∆F/F) of Seta and Leta during each motor patterns. While both Seta and Leta are active during BW, only Seta is active during AT and PT. \*\*\**p*0.001, ANOVA with post-hoc Tukey HSD test, compared to QS. (f-f”) ROI correlation mapping in *5-HT2A-GAL4*-targeted cells. Unlike in c, ROI mapping was applied only to *5-HT2A-GAL4*-targeted cells. The morphology of Seta (f, f”) and Leta (f’, f”) can be seen more clearly. Scale bar 100 *µ*m.

We next studied the morphology of Seta and Leta neurons. Since neurites of *5-HT2A-GAL4*-positive neurons extensively overlap, it was impossible to determine the morphology of individual neurons when markers such as mCD8-GFP were expressed in all GAL4-positive cells. We therefore first tried to reveal the morphology of Seta and Leta based on their activity by calculating the Pearson’s r. As shown in Fig. 4c”, f-f” and S4, this method identified voxels which are synchronously active and thus likely belong to each of the neurons, revealing the outline of their morphology. We then searched for these neurons in cells individually labeled by single-color and multi-color flp-out techniques^32^ and studied their morphology more in detail. The Seta neuron was found to be located in a ventro-medial region of T1 neuromere and extended its axon and dendrites in the dorsal neuropile (Fig. 5a, c). The axon of the neuron crossed the midline and then extended posteriorly along the CI tract (Fas2-coordinate^33^) to A1 neuromere. The dendrites extended posteriorly near the DM tract to T3 in the ipsilateral side. The Leta neuron was located in a medio-lateral region of T3 neuromere and extended its axon and dendrites in the dorsal neuropile (Fig. 5b, d). The axon of Leta neurons also crossed the midline and then bifurcated to extend posteriorly toward A6 neuromere and anteriorly toward T1 segment (Fig. 5b, d). The dendrites of Leta neurons innervated a medial region of the dorsal neuropil in T1-A3 neuromeres. Both Seta and Leta neurons were weakly stained with anti-Choline acetyltransferase (ChAT) antibodies (Fig. 5e, f). As will be described below, Seta and Leta also expressed other markers of cholinergic neurons, Cha-GAL80 and Cha3.3-GAL80. Thus, these neurons are likely cholinergic.

**Figure 5.**
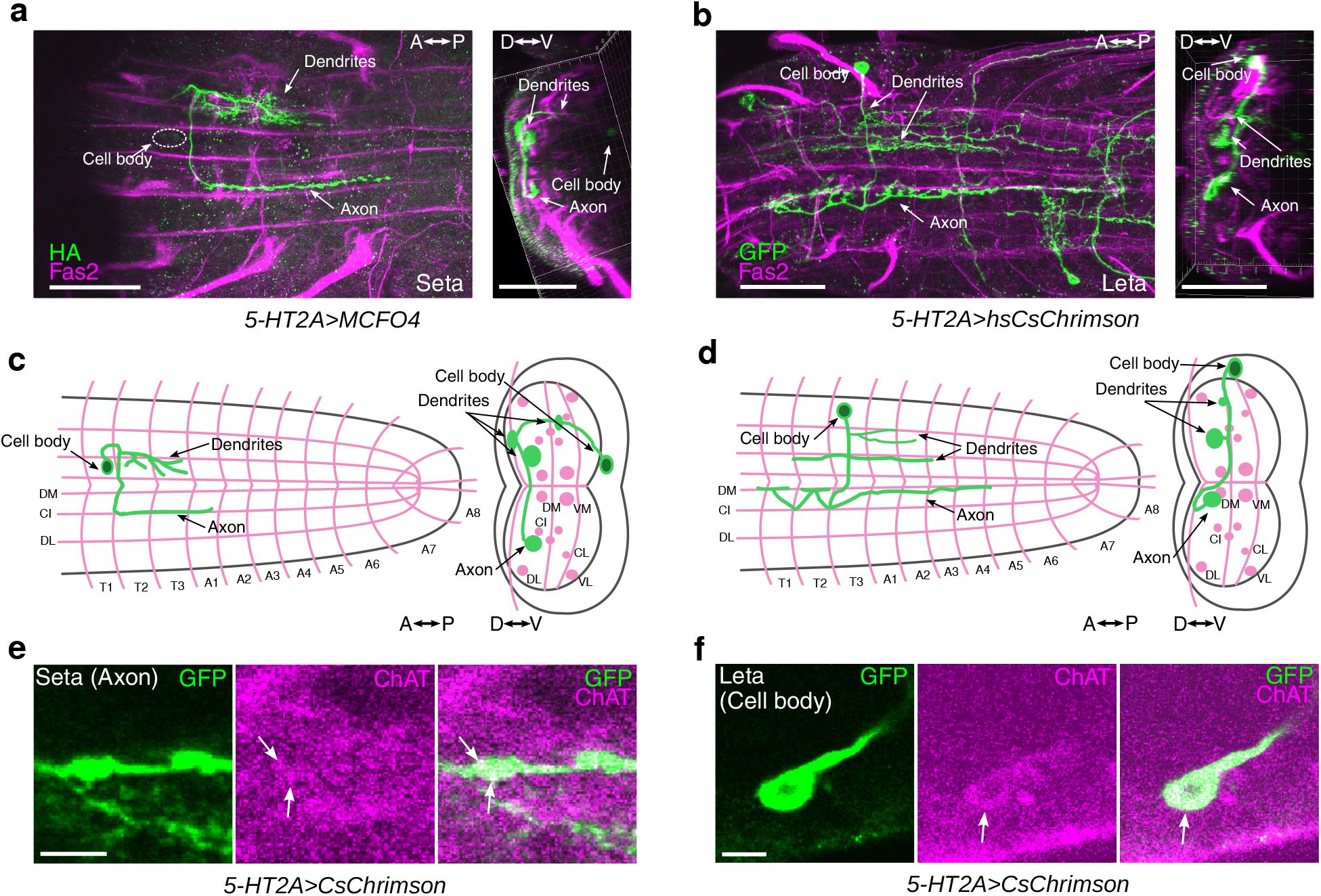
Morphology and neurotransmitter identity of Seta and Leta neurons. (a-b) Representative single neurons of Seta (a) and Leta (b) obtained by the MCFO method and hs-CsChrimson method in wandering third instar larvae, respectively (green). Fas2 expression is also shown (magenta). Dorsal (left) and cross-sectional (right) views are shown. (c-d) Schematic representation of the morphology of Seta (c) and Leta (d). Fas2 coordinates are shown as landmarks (magenta). (e-f) Co-staining for CsChrimson (expression driven by *5-HT2A-GAL4*, green) in Seta (e) and Leta (f) and for ChAT(magenta). A single plane focusing on the axon terminal of Seta (e) and cell body of Leta (f) is shown. Staining in different parts of the cells were shown because only the cell body and axons can be clearly identifiable as belonging to Leta and Seta, respectively, among the Gal4-targeted neurons. C: central, D: dorsal, I: intermediate, L: lateral, M: median, V: ventral, A: anterior, P: posterior. Scale bars, 50 *µ*m (a, b), 5 *µ*m (e, f). Note that b is identical to Fig. 6e.

### Optogenetic activation of Seta and Leta neurons increases backward locomotion

Since Seta and Leta are specifically active during backward but not forward fictive locomotion (Fig. 4d-e), they are potentially involved in the induction of backward locomotion (Fig. 6a). We therefore next asked if forced activation of these neurons is sufficient to induce backward peristalsis. We first studied the effects of activating all *5-HT2A-GAL4*-positive cells during locomotion. We expressed a red-shifted channelrhodopsin, CsChrimson^34^ and optogenetically activated these neurons in freely moving larvae. Light application significantly increased backward peristalsis (Fig. 6b), indicating that activation of the *5-HT2A* neurons promotes backward locomotion. To narrow down candidate neurons responsible for the optogenetic induction of backward locomotion, we used intersectional methods. *tsh-GAL80* and *cha3.3-GAL80* or *cha-GAL80*suppress the function of GAL4 in a majority of neurons in the VNC and cholinergic neurons, respectively. Accordingly, expression of CsChrimson driven by *5-HT2A-GAL4* was abolished in Seta and Leta neurons when *tsh-GAL80*, *cha3.3-GAL80* or *cha-GAL80* was combined with the GAL4 (Fig. S5). In these larvae, optogenetic induction of backward locomotion was significantly reduced (Fig. 6b). These results implicate cholinergic *5-HT2A* neurons in the VNC, including Seta and Leta, in the induction of backward locomotion. To further narrow down the expression of CsChrimson, we next used the flp-out technique to generate clones of cells expressing CsChrimson among the *5-HT2A-GAL4*-positive cells. We optimized the heat-shock condition so that less than ten neurons in the VNC expressed CsChrimson. We stimulated larvae with light and recorded the behavioral responses of 180 larvae upon optogenetic stimulation. The larvae were then dissected and the neurons expressing CsChrimson were identified in individual larvae. We observed a strong correlation between the expression of CsChrimson in Seta or Leta neurons and induction of backward locomotion (Fig. 6c-e). In contrast, such correlation was not seen for other *5-HT2A* neurons in the VNC (Fig. S6). These results indicate that activation of Seta or Leta promotes backward locomotion in the larvae.

**Figure 6.**
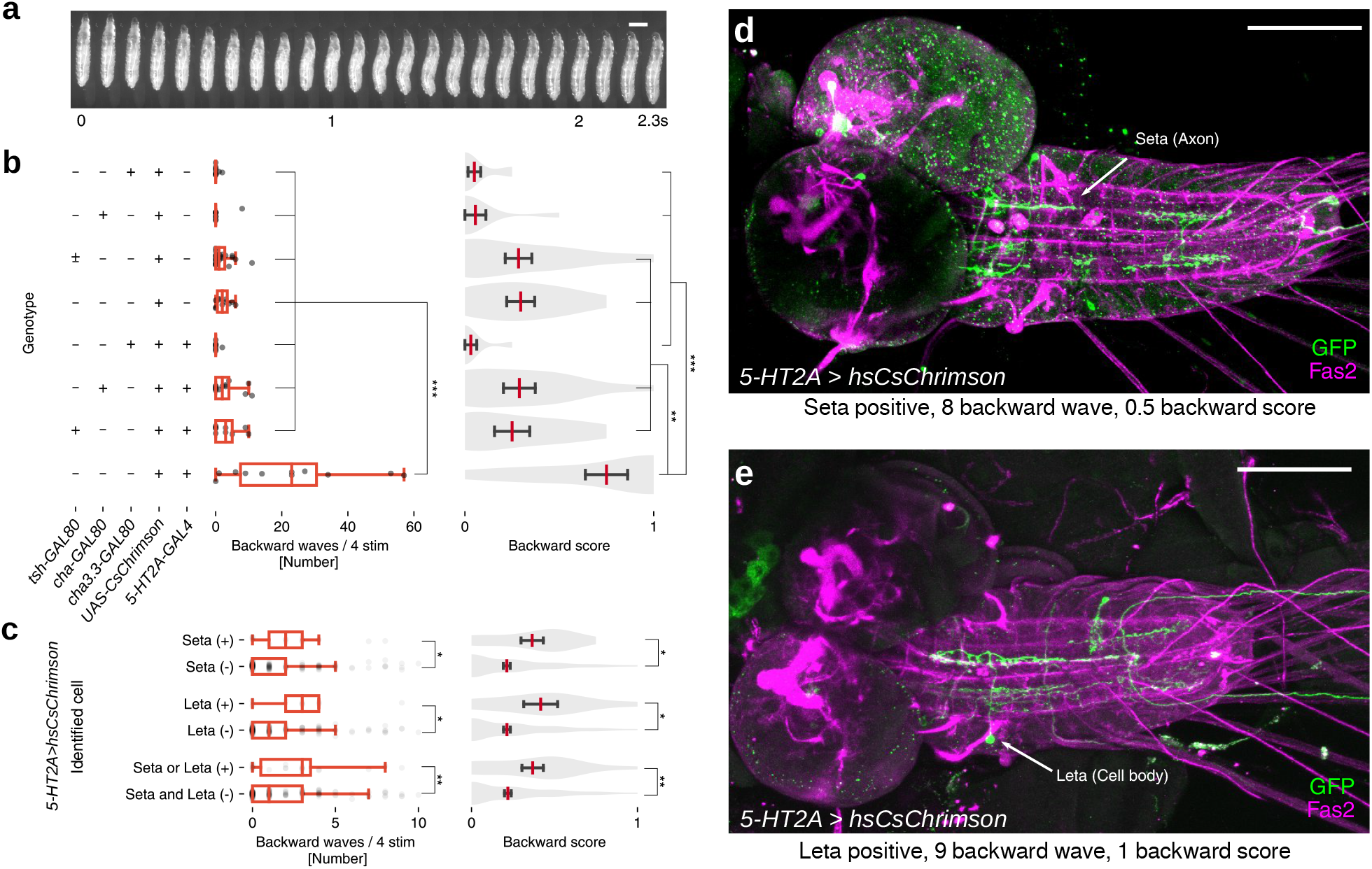
Optogenetic activation of the Seta or Leta neuron increases backward peristalsis. (a) Montage of a representative backward peristalsis of the 3rd instar larva. Scale bar 1mm. (b) Optogenetic activation of *5-HT2A-GAL4* targeted cells with or without *tsh-GAL80*, *cha3.3-GAL80*or *cha-GAL80*. Quantification of the number of backward peristalses (left, boxplot with scatter) and backward score (right, violinplot with mean and SEM) in each genotype is shown. n=^10, 9, 22, 11, 8, 13, 9^, 10 (from top to bottom). \**p*0.05, \*\**p*0.01, \*\*\**p*0.001, ANOVA with post-hoc Tukey HSD test. (c-e) Optogenetic activation of small subsets of *5-HT2A-GAL4* targeted cells. (c) Quantification of backward locomotion as in b in larvae with or without CsChrimson expression in Seta or Leta. \**p*0.05, \*\**p*0.01, Mann-Whitney *U*test. p=0.016 (waves), 0.010 (score) for Seta (n=13 (+), 167 (-)), p=0.015 (waves), 0.015 (score) for Leta (n=9 (+), 171(-)), p=0.005 (waves), 0.005 (score) for Seta or/and Leta (n=19 (+), 161 (-)). (d, e) Examples of CsChrimson expression. A larva with expression in a Seta neuron (d) and Leta neuron (d). The number of backward peristalsis induced and backward score is also shown. Scale bar 100 *µ*m.

### Larval serotonergic system is required for backward locomotion

We next tested if *5-HT2A-GAL4* neurons including Seta and Leta are required for the induction of backward peristalsis. We induced backward locomotion in free moving larvae by shining blue light, which is an aversive stimulus^35^. Thermogenetically inhibiting *5-HT2A-GAL4* cells using temperature-sensitive Shibire^36^ significantly decreased the induction of backward peristalsis (Fig. 7a) compared to control larvae, indicating that *5-HT2A* cells are required for proper induction of backward locomotion. Since *5-HT2A* expressing neurons are likely to be in part regulated by 5-HT, we next asked if 5-HT releasing neurons are also involved in the induction of backward locomotion. We used the same light-induced behavioral assay and found that thermogenetic inhibition of 5-HT neurons with Shibire^*ts*^ (whose expression driven by *trh-GAL4*, which specifically targets 5-HT neurons^37^) significantly reduced the rate of backward locomotion (Fig. 7b). These results suggest the possibility that 5-HT neurons induce backward locomotion by modulating *5-HT2A* neurons, including Seta and Leta, via the 5-HT2A receptor. To test this, we next studied the effect of Ketanserin, a specific antagonist for the 5-HT2 receptor^38^ on fictive backward locomotion. A recent study reported that Ketanserin has no effect on 5-HT2B^39^, the only other 5-HT2 receptor in *Drosophila*, suggesting that the drug is potentially specific to 5-HT2A receptors. We used our methodology described above to quantify the occurrence of backward locomotion before and after the addition of Ketanserin. Frequency of motor patterns related to backward locomotion, AT and BW, were significantly reduced upon application of the drug (Fig. 7c, see S7a-b for details). In contrast, there was no change in the frequency of FW. These results render further support for the notion that 5-HT modulates backward locomotion via the 5-HT2A receptor and neurons. We also studied a loss-of-function mutation of *5-HT2A* (*5-HT2APL*^40^) but found no effect on backward locomotion (Fig. 7b). This could be due to the fact that the mutation is partial loss-of-function^40^ and/or compensations during development. Finally, we studied the activity of 5-HT neurons during fictive locomotion. Since they are involved in the regulation of backward locomotion, they may be selectively active during the relevant motor patterns. We expressed GCaMP6f in 5-HT neurons using *trh-GAL4* as a driver and studied their activity during fictive locomotion in isolated CNSs (Fig. 7d). We again used our methodology to classify motor patterns seen in the activity of 5-HT cells. As when GCaMP6f was expressed in motor neurons or all neurons, the four motor patterns were extracted (Fig. 7e, f). This indicates that 5-HT neurons in the VNC show activities related to all fictive motor patterns, including the segmental propagation corresponding to forward and backward locomotion. However, the intensity of the activity was much higher during motor patterns related to backward locomotion, AT and BW, compared to those related to forward locomotion, PT and FW (Fig. 7g). These results suggest that 5-HT is released preferentially during backward locomotion.

**Figure 7.**
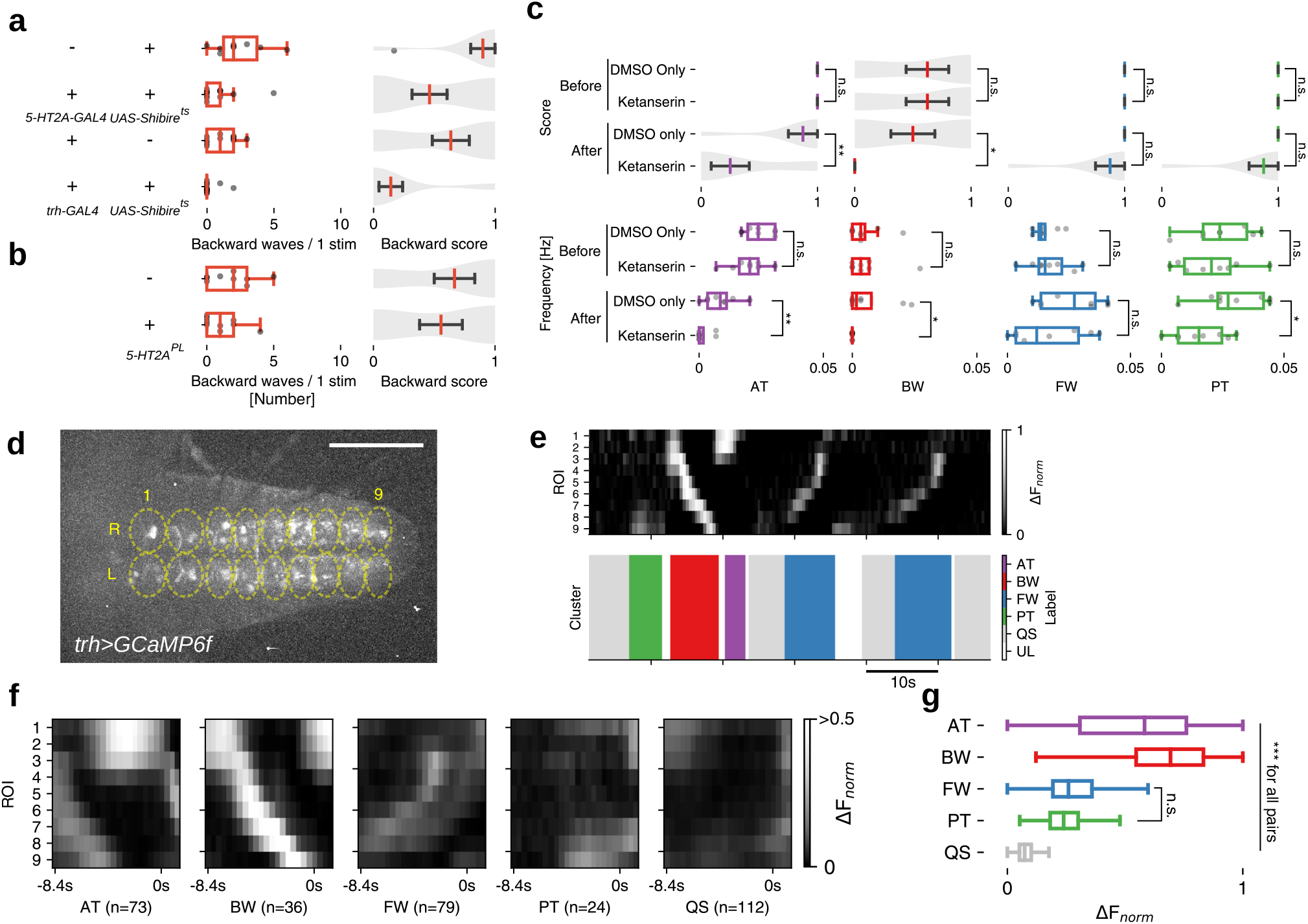
Implication of the serotonergic system in backward locomotion. (a) *5-HT2A-GAL4* and serotonergic neurons are required for proper backward locomotion. Quantification of backward locomotion as in Fig. 6b when *5-HT2A-GAL4* or *trh-GAL4* targeted neurons are thermogenetically inhibited (n=10, 13, 11 and 14 from top to bottom). p=0.004 (waves), 0.016 (score) for *5-HT2A>Shibire^t s^*, p=0.005 (waves), 0.006 (score) for *trh>Shibire^t s^*. (b) 5-HT2A homozygous mutants (*5-HT2APL*) did not show significant difference in backward locomotion. n=9, each group. p=0.19 (waves), 0.32 (score) for *5-HT2A^PL^*. \**p*0.05, \*\**p*0.01, Mann-Whitney *U* test. (c)A 5-HT2 antagonist, Ketanserin, reduces fictive backward locomotion. Motor patterns were extracted from activity data of *5-HT2A, RRa>GCaMP6f*. p=0.001 (waves), 0.002 (score) for AT, p=0.023 (waves) 0.016 (score) for BW, p=0.15 (waves), 0.02 (score) for PT, otherwise p*>*0.05. (d-g) Calcium imaging of serotonergic neurons. (d) Neural activity in *trh>GCaMP6f* larvae was captured using 18 ROIs. (e) Extracted motor patterns. Preprocessed fluorescence pattern (top) and assigned motor patterns (bottom). (f) Mean activity pattern aligned with the end point of each motor pattern. Average duration of each motor pattern were AT:2.2 ±0.2s, BW:2.6 ±0.1s, FW:1.8 ±0.1s, PT:1.5 ±0.1s, QS:9.1 ±0.8s, (mean±standard deviation). (g) Maximum fluorescence change during each motor patterns. \*\*\**p*0.001, ANOVA with post-hoc Tukey HSD test. Scale bar 100 *µ*m. See also Fig. S7.

## Discussion

Classifying activity patterns is crucial in studying large-scale calcium imaging data. Previous classifying methods often require manual optimization thereby making data analysis laborious and less flexible. Recently, rapid development in CNN has produced remarkable results in various fields including biomedical research^41^. Given the fact that human can perceive the neuronal activity pattern as images resembling patterns of MNIST handwritten digits^24^, we thought it would be possible to distinguish neuronal activity patterns using a CNN pre-trained by image classification. As expected, we found that motor activity patterns could be classified by the deep convolutional feature extracted from Conv4 3 of VGG-16 model and HAC. The VGG-16 model^28^ was used since it has a relatively simple architecture among the contemporary CNN models^25, 42^ and has been broadly used for transfer learning. Since manual optimization is minimized, our method is less laborious compared to previous methods and could easily be adopted to imaging data obtained under various imaging conditions. Our method is also more objective since the unsupervised clustering reduces human interpretation. Furthermore, while our method was only applied to the larval preparation in this study, similar strategies would be useful for classifying activity data in other systems. Although our method was only performed as offline processing to fit HAC on the data and label its clusters, it may be performed as online processing since HAC can predict motor patterns in unseen data once it is fitted during the offline processing. Moreover, the performance of our method may be improved by introducing supervised learning such as fully connected neural networks and random forest classifier (transfer learning). Our methodology will greatly accelerate future analyses of fictive locomotion in *Drosophila* larvae.

The methodology successfully extracted all of the four major motor patterns, FW, BW, AT and PT, which have previously been detected by human inspection in an isolated nerve cord^13^. Furthermore, it did so in activity data of different cells: two pairs of MNs in each neuromere (aCC and RP2, Fig. 2), all neurons (Fig. 3) and small subsets of interneurons (5-HT neurons, Fig. 7), the data of which were acquired under various time resolution (180-420ms per volume, Fig. 2-4,7). It also works with small sample size (3-12 samples, Fig. 2-4, 7). Thus, our method appears to be robust and works with various numbers and types of neurons and under different imaging conditions.

We analyzed transition probabilities between the four motor patterns and found that they are biased as observed in intact larvae. This is consistent with that the behavioral decision of the intact larva is biased^30, 31, 43^ and suggests that the bias partly derives from the intrinsic properties of the CPGs. However, while free moving larvae on a substrate rarely show backward locomotion, backward waves occur frequently in the isolated CNS. The Markov chain found in the isolated CNS may be more closely related to larval behaviors within the food^44^ or when sensory input is genetically removed^45^, where backward locomotion is more frequently seen.

We also used correlation mapping to identify voxels (Fig. 3a-c, 4c-d, 7e) and cells (Fig. 4d-e) showing correlated activity with each of the motor patterns classified by VGG-16 and HAC. Taken together, these methodologies establish a pipeline that efficiently extracts motor patterns and the associated cells from the calcium imaging data. We first applied the pipeline to activity data derived from all neurons (*elav>GCaMP6f*) and succeeded in identifying cellular structures which are preferentially active during PT and BW. Those correlated with PT are longitudinal axons that run across the VNC (Fig. 3c””). Since the voxel-wise correlation mapping was limited by imaging resolution and therefore did not allow detection of the cell body, it is currently unknown whether the axons belong to ascending or descending neurons. Those correlated with BW are neurites spanning a few anterior neuromeres. To our knowledge, neurons with such morphologies have not been identified to be associated with PT or BW. Future identification of sparse GAL4 lines targeting these neurons will enable functional analyses of the roles of these neurons in motor regulation.

Although we could identify some cellular structures specific to PT and BW, most of the voxels which were preferentially active for each of the motor patterns were intermingled with each other and thus were difficult to be individually identified. We therefore next applied the pipeline to neurons targeted by sparse GAL4 lines. As candidates, we focused on interneurons that express NM receptors since NMs are known to regulate various animal behaviors such as locomotion, foraging, feeding, aggression and courtship^46–48^ Since antibodies are not available, previous studies used GAL4 lines (either enhancer-trap insertions or driven by cis-elements) which are expected to mimic the endogenous gene regulation to study the expression of some of the NM-receptors^49^. However, for most of the NM-receptors, GAL4 lines have not been available and their expression pattern was therefore completely unknown. In this study, we systematically generated GAL4 lines targeting most of the NM receptors in *Drosophila* using CRISPR/Cas9-mediated gene knock-in. Since GAL4 was inserted in frame with the endogenous proteins, the GAL4 lines are expected to reproduce the expression of the NM-receptors in a more precise manner than previously reported GAL4 lines. We found that most of the NM-receptor GAL4 lines drive expression broadly in large subsets of neurons in the CNS, sensory neurons and muscles (Fig. S3). However, one of the lines *5-HT2A-GAL4* showed sparse expression and application of the pipeline to this GAL4 line allowed us to identify two interneurons termed Seta and Leta correlated with backward locomotion. Subsequent optogenetic experiments showed that activation of these neurons increases the frequency of fictive backward locomotion (Fig. 6). Furthermore, thermogenetic experiments showed that backward locomotion of the larvae induced by aversive stimuli is reduced when *5-HT2A-GAL4*-targeted neurons are inhibited (Fig. 7a). The involvement of *5-HT2A* neurons in the regulation of backward locomotion was further supported by the effects of Ketanserin, antagonist putatively specific to 5-HT2A receptor, on fictive backward locomotion (Fig. 7b). While these loss-of-function experiments do not specifically target Seta and Leta, the results of optogenetic activation and the fact that these are the only cells we detected as active during BW strongly suggest that these neurons are required for proper execution of backward locomotion.

We also studied the role of neurons expressing the ligand of the 5-HT2A receptor, Serotonin (5-HT). 5-HT is a well-known NM that plays essential roles in the regulation or modulation of animal behaviors. For instance, 5-HT has been implicated in the regulation of escape behavior and crawling in the mollusk *Tritonia*^50^, dwelling, slowing and transition from crawling to swimming in the nematode *Caenorhabditis elegans*^51–53^ and nociception and turning in larval *Drosophila*^35, 54^. We found that thermogenetic inhibition of 5-HT neurons reduced the frequency of backward locomotion in the larvae, as when *5-HT2A* cells are inhibited. We also showed that 5-HT neurons are more strongly activated during backward fictive locomotion than during forward locomotion. Furthermore, effects of Ketanserin indicate that 5-HT regulate backward locomotion in part via *5-HT2A* cells. Taken together, our results strongly suggest that 5-HT modulates backward locomotion by acting on Seta and Leta neurons via the 5-HT2A receptor.

It is clear that analysis of neuronal activity at the system level is crucial for understanding the circuit underpinnings of CPGs^8, 12, 55^. Accordingly, improved methodologies and pipelines that link large-scale activity data to motor patterns and associated component neurons would greatly facilitate future studies of CPGs. The methodology and pipeline we developed in this study are based on open source and can be easily handled. They allow rapid and robust classification of motor patterns under various imaging conditions. We showed the utility of the pipeline by applying it to *5-HT2A-GAL4* neurons and successfully identifying interneurons involved in the regulation of backward locomotion. In the future, application of the pipeline to other GAL4 lines would enable systematic search of interneurons involved in motor regulation. This work lays the foundation for the quantitative analysis of neuronal activity in larval Drosophila and shows how state-of-art machine learning technique can be applied to biological research.

## Methods

### Fly strains

The following fly strains were used: *yw*, *10xUAS-IVS-mCD8::GFP*^56^, *UAS-GCaMP6f*^57^, *cha-GAL80*^36^, *cha3.3-GAL80*^36^, *RRa-GAL4*^58^, *tsh-GAL80*^59^, *UAS-CsChrimson-mVenus*^34^, *UAS-Shibiret s*^36^, *elav-GAL4*^60^, *UAS-FRT-stop-FRT-CsChrimson-mVenus*^20^,*UAS-MCFO4* 32. *w∗*; *P(FRT(w^hs^))2A P(neoFRT)82B PBac(GAL4D,EYFP)5-HT2APL*^00052^ (Bloomington Drosophila Stock Center (BDSC) stock No. 19367) was used as an allele of *5-HT2A*(*5-HT2A^PL^*^00052^ ^40^). *w*^1118^; *P(FRT(w^hs^))2A P(neoFRT)82B* (BDSC stock No. 8218) was used as a control for the *5-HT2A*mutant. Flies were raised on conventional cornmeal agar medium at 25◦C except for those expressing Shibire*^ts^*which were raised at 18◦C.

### Generation of NM receptor GAL4 lines

GAL4 knock-ins were generated by targeted transgene insertion using the CRISPR/Cas9 system (Kondo and Ueda 2013). A transgene encoding the 2A peptide and the GAL4 transcription factor was inserted in front of the stop codon of each NM receptor gene. The translated product, which is a fusion protein of the NM receptor, the 2A peptide and GAL^4^, is cleaved at the 2A peptide so that both the NM receptor and GAL4 are expressed in the cell. Lines were generated for the following genes: *CG9652 (Dop1R1)*, *CG33517 (Dop2R)*, *CG18314 (DopEcR)*, *CG15113 (5-HT1B)*, *CG1056 (5-HT2A)*, *CG42796 (5-HT2B)*, *CG120732 (5-HT7)*, *CG33976 (Octb2R)*, *CG42244 (Octb3R)*, *CG3856 (Oamb)*, *CG16766 (TyrRII)*.

### Optogenetic activation

Hatched larvae were raised on regular food at 25^◦^C until second instar (∼72 hours after egg laying; hAEL) and transferred to an agar plate with yeast paste containing 1mM all-trans retinal 24 hours before optogenetic experiments. All optogenetic experiments were done at 25◦C under dim light. Third instar larvae (∼96 hAEL) were washed in deionized water to remove residual yeast paste and acclimated on an agar plate for a minute before activation experiments. Larval locomotion on the agar plate was recorded using a stereoscopic microscope (SZX16 or MVX10; Olympus) and a CCD camera (XCD-V60, Sony) at 15 frames per second. Free-moving larvae were stimulated with red light (660nm, LED, Thorlabs) at 1mW/mm^2^ for 20 seconds four times with 20 seconds interval. Number of backward peristalses was counted manually. We quantified backward locomotion in two ways: ”Backward wave” is total number of backward peristalsis in four stimulations, ”backward score” is defined as 0 (no backward peristalsis) or 0.25 (one and more than one backward peristalsis) per stimulus so that the maximum score is 1 for four stimuli in one trial. Backward waves and backward score measure how strong the phenotype is and how constant it is, respectively. We used progeny larvae from the cross between *yw* and a UAS effector line as a control.

### Stochastic optogenetic activation of *5-HT2A-GAL4* targeted neurons using Flp-out technique

Female *20XUAS>dsFRT>CsChrimson::mVenus (attP18), pBPhsFlp2::Pest (attP3)* flies were crossed with *5-HT2A-GAL4* males. Eggs were collected for 24 hours on an apple-juice agar plate with yeast paste. The eggs were raised at 25◦C for 24 hours. Then, heat shock (37◦C) was applied for 45 min to express Flippase. After that, the larvae were transferred to another plate, on which 1mM All-trans-Retinal-containing yeast paste was spread, and raised at 25◦C for 48-72 hours. Optogenetic experiments were done as described above. 180 animals were tested, and the expression patterns of CsChrimson::mVenus in all larvae were examined by immunostaining. Based on the morphology, we labeled a subset of the neurons: Seta, Leta, TLN (Thoracic Long-axon Neuron), TUN (Thoracic U-shaped Neuron), and TBN (Thoracic Bump Neuron) (Fig. S6). We selected these neurons for the following reasons: located in thoracic neuromeres, not segmentally repeated and easy to identify by their characteristic morphology.

### Temperature shift experiments

Parental flies and eggs were raised at 22◦C for Shibire*^ts^* experiments. For blocking neural activity, larvae were put on an agar plate held at 32◦C on a heat plate (Thermo plate, Tokai Hit, Japan). To induce photo-avoidance backward locomotion, we stimulated free-moving larvae with blue light (488nm, mercury lamp) at 0.5mW/mm2. Locomotion of larvae was recorded and analyzed as described in the optogenetic activation section. Progeny larvae from *yw* a crossed between UAS effector line and *trh-GAL4* or *5-HT2A-GAL4* were used as a control for *5-HT2A>Shibire^ts^* or *trh>Shibire^ts^*, respectively.

### Immunohistochemistry

Wandering third instar larvae were dissected in phosphate buffered saline (PBS) and fixed with 4% formaldehyde in PBS for 30 min at room temperature. Fixed larvae were washed with 0.2% Triton X-100 in PBS (hereafter PBT) and blocked with 5% normal goat serum (NGS). Blocked larvae were incubated at 4◦C with the primary antibodies over 12 hours. The larvae were then washed with PBT and incubated at 4◦C with the secondary antibodies over 12 hours. We used confocal scanning microscope (Fluoview FV1000, Olympus) with a water immersion objective 20x and 60x lenses for fluorescence imaging. The frame size of the images varied from 1024×768 pixels to 1600×1200 pixels. The images were processed and analyzed using Fiji (ImageJ) and Imaris (Bitplane) software. The following primary antibodies were used: rabbit anti-GFP (Af2020, 1:1000, Frontier Institute), mouse anti-Fas2 (1D4, 1:10, DSHB), rabbit anti-HA (C29F4, 1:1500, Cell Signaling Technology), mouse anti-ChAT (4B1, 1:5, DSHB). The following secondary antibodies were used: goat anti-mouse Cy5, goat anti-rabbit Alexa Fluor 488 (Life Technologies).

### Calcium Imaging

We selected wandering third instar larvae (96-120hAEL) and isolated the CNS from the bodywall. The larval CNS was placed dorsal-side up on a MAS-coated slide glass (cell adhesive slide glass, Matsunami glass, JAPAN) and covered with TES buffer (TES 5 mM, NaCl 135 mM, KCl 5 mM, CaCl_2_ 2 mM, MgCl_2_ 4mM, Sucrose 36 mM). Imaging was performed with a spinning-disk confocal unit (CSU21, Yokogawa) and EMCCD camera (iXon, Andor) installed on Axioskop2 FS microscope (Zeiss). 20x water-immersion objective lense was used. Imaging conditions for each genotype are described in Tab. 1.

**Table 1.** Imaging conditions for each genotype.

| Genotype | imaging time per volume | z | total frames | sample No. |
| --- | --- | --- | --- | --- |
| *RRa>GCaMP6f* | 180 ms | 3 | 2048 | 12 |
| *elav>GCaMP6f* | 180 ms | 3 | 1500 | 3 |
| *5-HT2A,RRa>GCaMP6f* | 420 ms | 2 | 2048 | 3 |
| *trh>GCaMP6f* | 420 ms | 2 | 2048 | 4 |

### Preprocessing for feature extraction

Changes in fluorescence intensity (F(t)) within 18 ROIs, which roughly correspond to hemineuromeres T2 to A7 at both sides, were measured. For each time point t (=0, 1, 2…) and each ROI, we defined the baseline (F*_base_* (t)) as the mean value of F(t-16) to F(t-1). Considering that action potential occurs when the GCaMP6f signal is rising^57^, and to minimize the effects of fluorescence degeneration, changes in fluorescence (∆F/F) were defined as (F(t)/F*_base_* (t)-1) *H*(F(t)/F*_base_* (t)-1) where *H*(*x*) is the Heaviside function so that the value of F(t)/F*_base_* (t)-1 less than 0 was treated as 0. Therefore, ∆F/F we defined indicates how much calcium signal is rising. The value of ∆F/F of ROIs from aCC motor neurons (*RRa>GCaMP6f*; *RRa,5-HT2A>GCaMP6f*) more than 2 was treated as 2 for value fitting because it occasionally becomes extremely high for unknown reasons. For image conversion, ∆F/F is scaled to between 0 to 1 using min-max transform (∆F*_norm_*) and multiplied by 255 and then converted to 8 bit integer. 18 ROIs are compressed to 9 ROIs using the maximum value among the values of left and right side and the 9-dimensional values were used for the following analyses.

### Feature extraction and clustering

We defined window size as 8 time frames (from t-4 to t+3) so that each window has 9 ∆F *norm* of ROIs and 8 time frames. Each window was transformed to a 72 pixel (resized from 9 ROIs) x 72 pixel (resized from 8 time frames) x 3 channel image where all channels have the same values. The deep convolutional features of the images were extracted from intermediate layer (Conv4 3) of ImageNet pre-trained VGG-16 model^28^. The features were compressed using global average pooling and clustered by hierarchical agglomerative clustering (Ward’s method, Tab. 2). Labeling of the clusters was performed by comparing average images of the first 20-30 clusters. The images showing high intensity in T2-A4 and A4-A7 are categorized as AT and PT, respectively. The images showing intensity flow from the posterior to anterior and from the anterior to posterior are categorized as FW and BW, respectively. The images showing nearly no intensity change are categorized as QS. We counted consecutive frames of the same label as one activity. The codes and datasets with ground truth used in Fig. 1-2 are available (https://github.com/Jeonghyuk/behavior-class.git).

**Table 2.** An example of data flow of sample with 2048 time frames.

| Analysis steps | Data shape | Detail |
| --- | --- | --- |
| ROIs | (18, 2048) | T2-A7, left and right |
| Smoothing and MinMax | (18, 2032) | Value 0-1 |
| Max | (9, 2032) | Represents segmental maximum activity |
| Window | (9, 8, 2024) | 9 ROIs x 8 frames x 2024 window |
| Image converting | (72, 72, 3, 2024) | 72 pixel x 72 pixel RGB image |
| Feature extraction | (512, 9, 9, 2024) | features of Conv4.3 |
| Global average pooling | (512, 2024) | for reduce parameter size |
| Hierarchical agglomerative clustering | (2024) | Ward's method |
| Label cluster data | (2032) | Compensation for 4 frames for front and rear |

### Correlation mapping

Registration of images was performed by ImageJ (NIH) and the Template matching plugin. For *RRa>GCaMP6f*, *5-HT2A,RRa>GCaMP6f* and *trh>GCaMP6f*, z stacked images were used for registration over time. For *elav>GCaMP6f*, each z plane was used for registration. Changes in fluorescence (∆F/F) were defined as described above. We defined a binary label vector as (1 for label, 0 for not label). For voxel-wise behavior correlation mapping (behavior mapping), Pearson’s r of ∆F/F of voxels and the binary label vector were calculated. For voxel-wise ROI correlation mapping (ROI mapping), Pearson’s r of ∆F/F of voxels and ∆F/F of ROIs were calculated. F(t) of a ROI is derived from the sum of F(t) of voxels whose Pearson’s correlation coefficient is 0.05 or higher.

### Bath application

Ketanserin (+)-tartrate salt (D006-10MG) was purchased from MERCK and stored at 4◦C. Ketanserin (10mM concentration) was solved in DMSO just before the experiment and stored at 4◦C. Experiments were performed with final concentration of Ketanserin in the bath to be 100 *µ*M. At this concentration, the effects of 5-HT on larval heart rate is known to be completely blocked^61^.

### Statistical analysis

We used Mann-Whitney *U* test and one-way analysis of variance (ANOVA) followed by Tukey’s HSD test for multiple comparisons. Statistical significance is denoted by asterisks: ***p0.001; **p0.01; *p0.05; n.s., not significant. All statistical tests were performed using Python. The results are stated as mean±s.d. or mean±SEM.

### Data Availability

The datasets generated during and/or analysed during the current study are available from the corresponding author on reasonable request.

## Acknowledgements

We are indebted to Seungho Lee for his excellent advice in applying neural network models. We also thank Youngtaek Yoon for help in calcium imaging. We are grateful to Bloomington Drosophila stock center and Kyoto stock center for reagents.

## Author contributions statement

J.P., H.K. and A.N. conceived and designed the experiments. J.P. performed the experiments and analyzed the data. S.K., H.T. and H.K. contributed reagents. J.P., H.K. and A.N. wrote the main manuscript. All authors reviewed the manuscript.

## Additional information

**Competing interests** The authors declare no competing interests.

---

**Figure S1.**
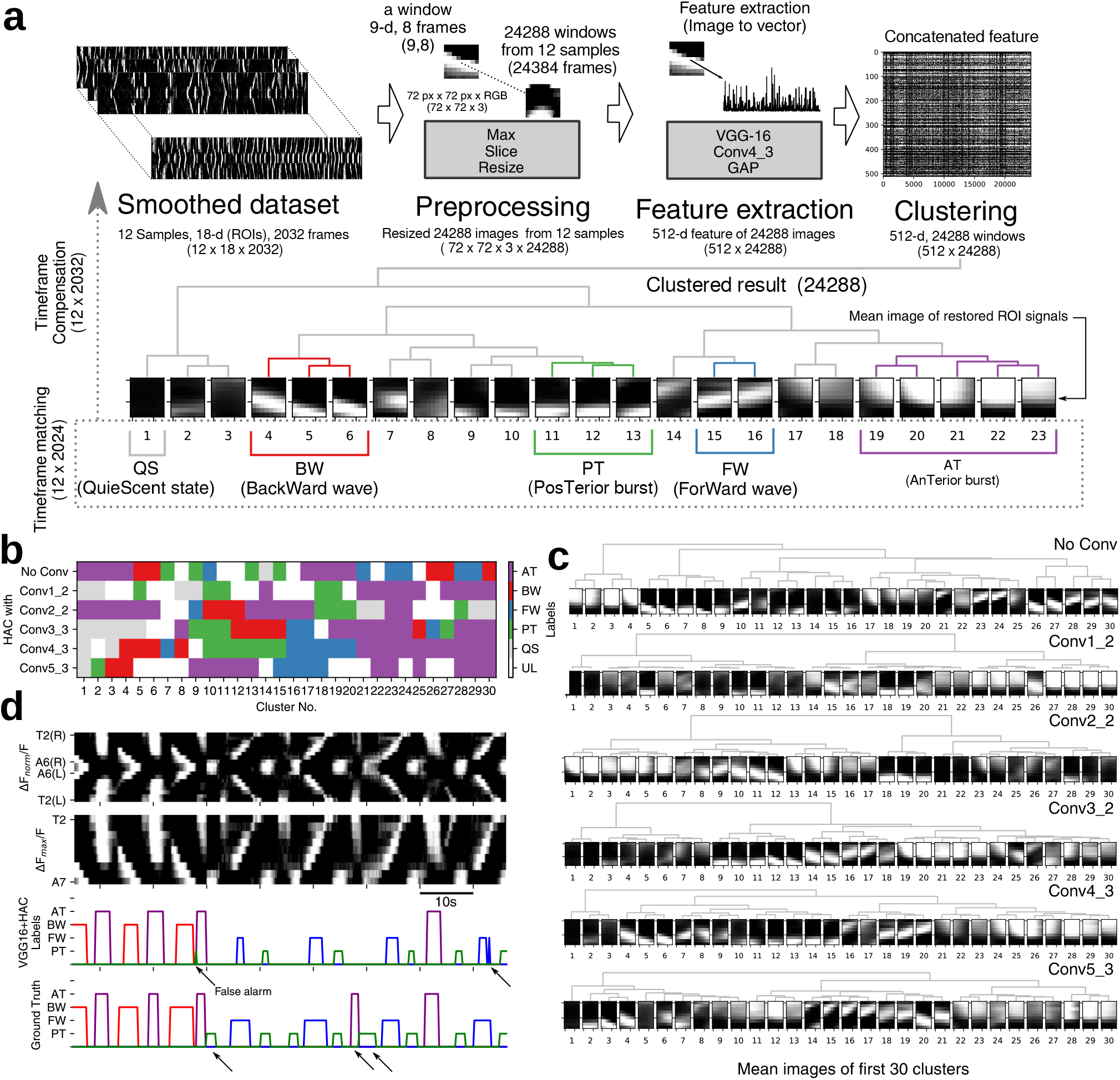
related to Fig. 1 and Methods. (a) Details of feature extraction and clustering. (b, c) Evaluation of cluster analysis using features from different layers of VGG16. Cluster analyses of the five convolution layers of VGG16 (Conv1 2, Conv2 2, Conv3 2, Conv4 3 and Conv5 3) and the original data. Note that input of HAC must be 1-d vector, windows are flattened to vectors for HAC without convolution layer (No Conv). Each cluster was manually inspected to determine if it represents a motor pattern. Conv4 3+HAC gave the best performance among the layers in that it identified all four motor patterns (AT, PT, FW and BW) and in a well-separated manner. (d) An example of comparison between the original activity data (top, 18-d ROI and 9-d ROI images), motor patterns assigned by VGG16+HAC (VGG16+HAC labels) and human inspection (ground truth). Visual cues indicate false alarms.

**Figure S2.**
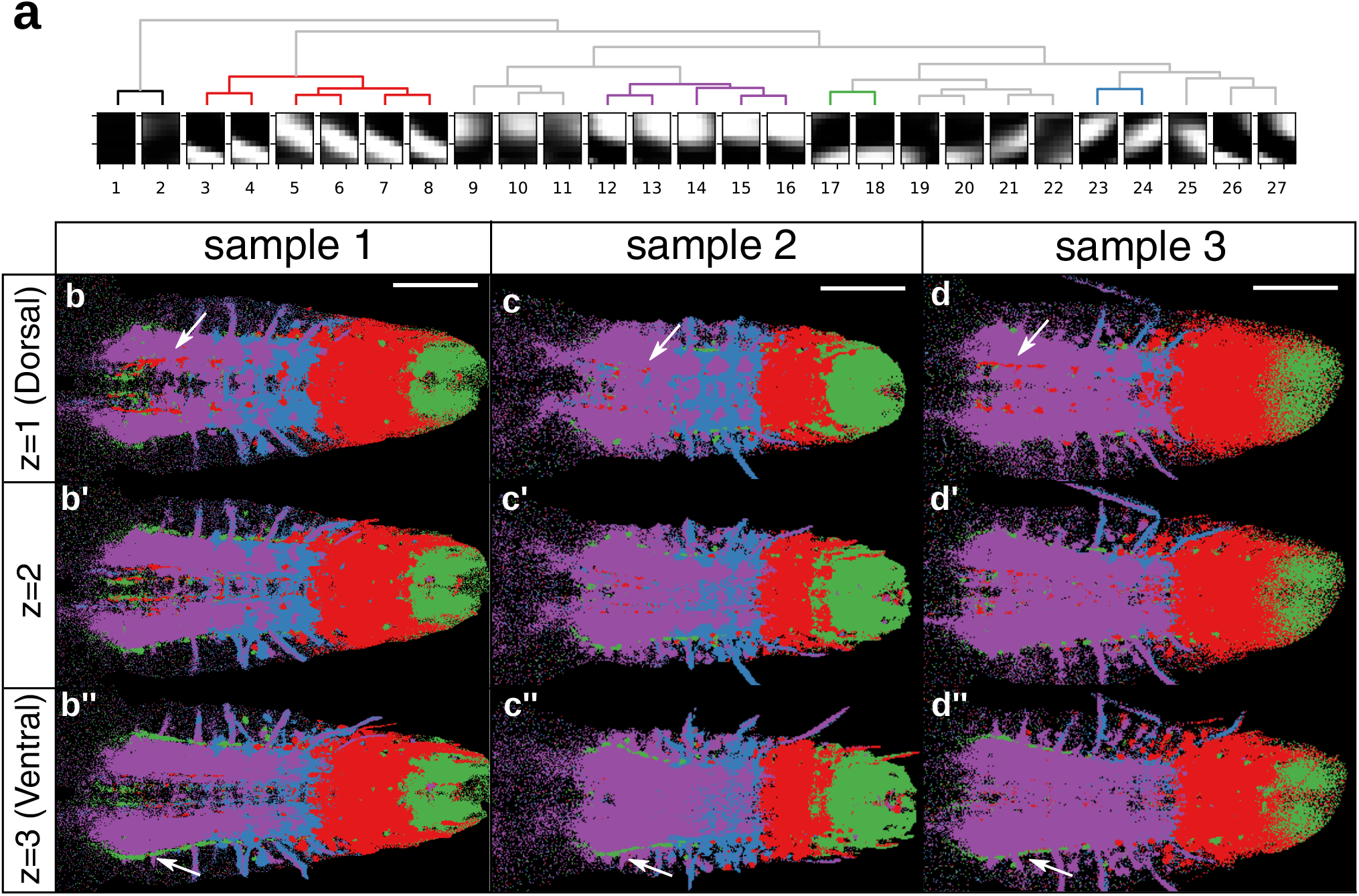
related to Fig. 3. (a) Clustering analyses of the Conv4 3 of VGG16 derived from activity data of all neurons. (b-d”) Dominant motor pattern mapping. Same as Fig. 3c but voxels are colored only for the most dominant motor pattern (highest Pearson’s r). Three z sections from dorsal to ventral are shown. Visual cues indicate structures specifically active during BW and PT. Note that b-b” is derived from the same sample as Fig. 3c. Scale bar 100 *µ*m

**Figure S3.**
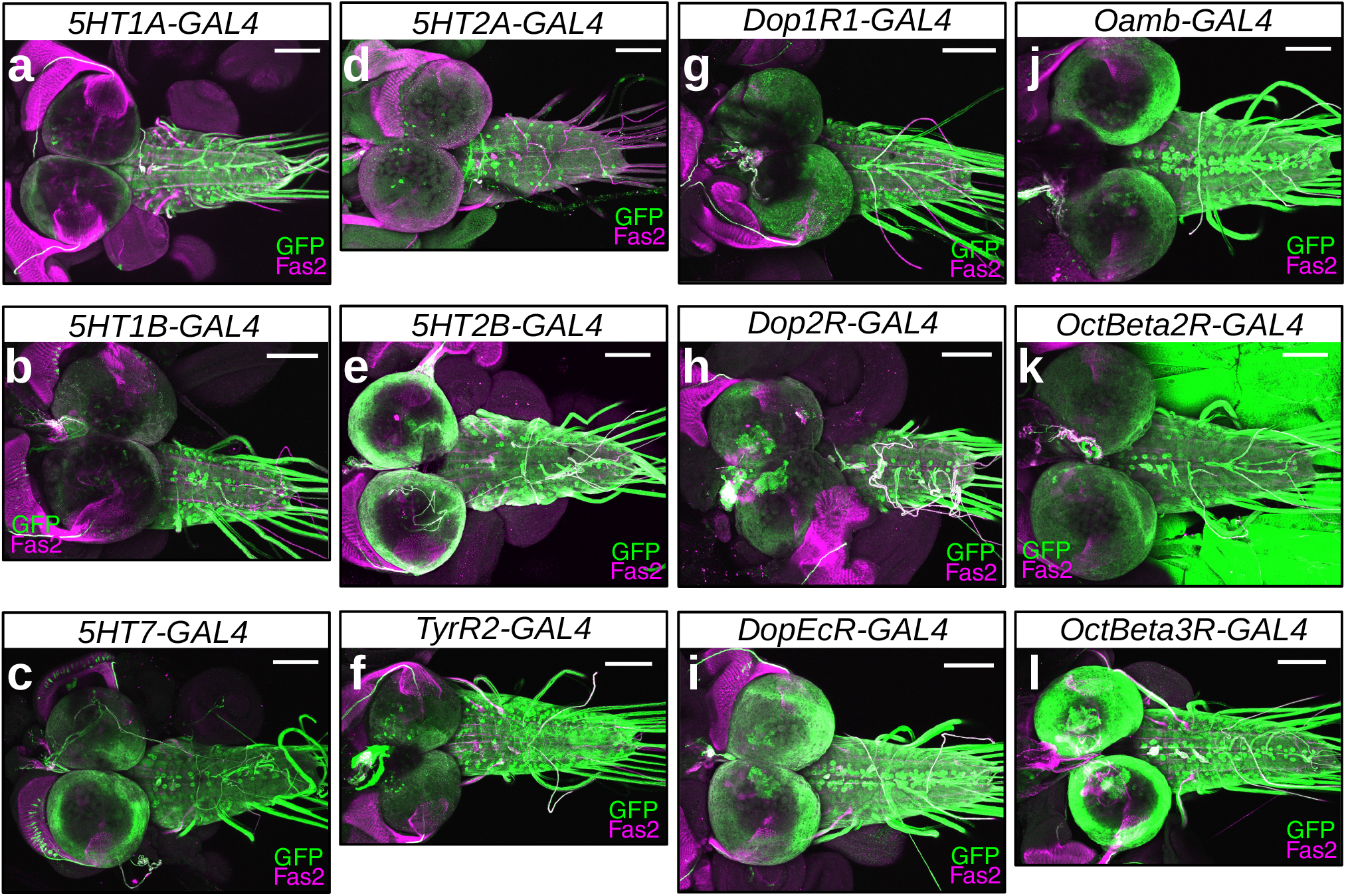
Expression of neuromodulator receptor GAL4 lines. related to Fig. 4. Fas2 (magenta) was used as a reference marker. *5-HT1A* (a), *5-HT1B* (b), *5-HT7* (c), *5-HT2A* (d), *5-HT2B* (e), *TyrR2* (f), *Dop1R1* (g), *Dop2R* (h), *DopEcR* (i), *Oamb* (j), *OctBeta2R* (k), *OctBeta3R* (l). Scale bar 100 *µ*m.

**Figure S4.**
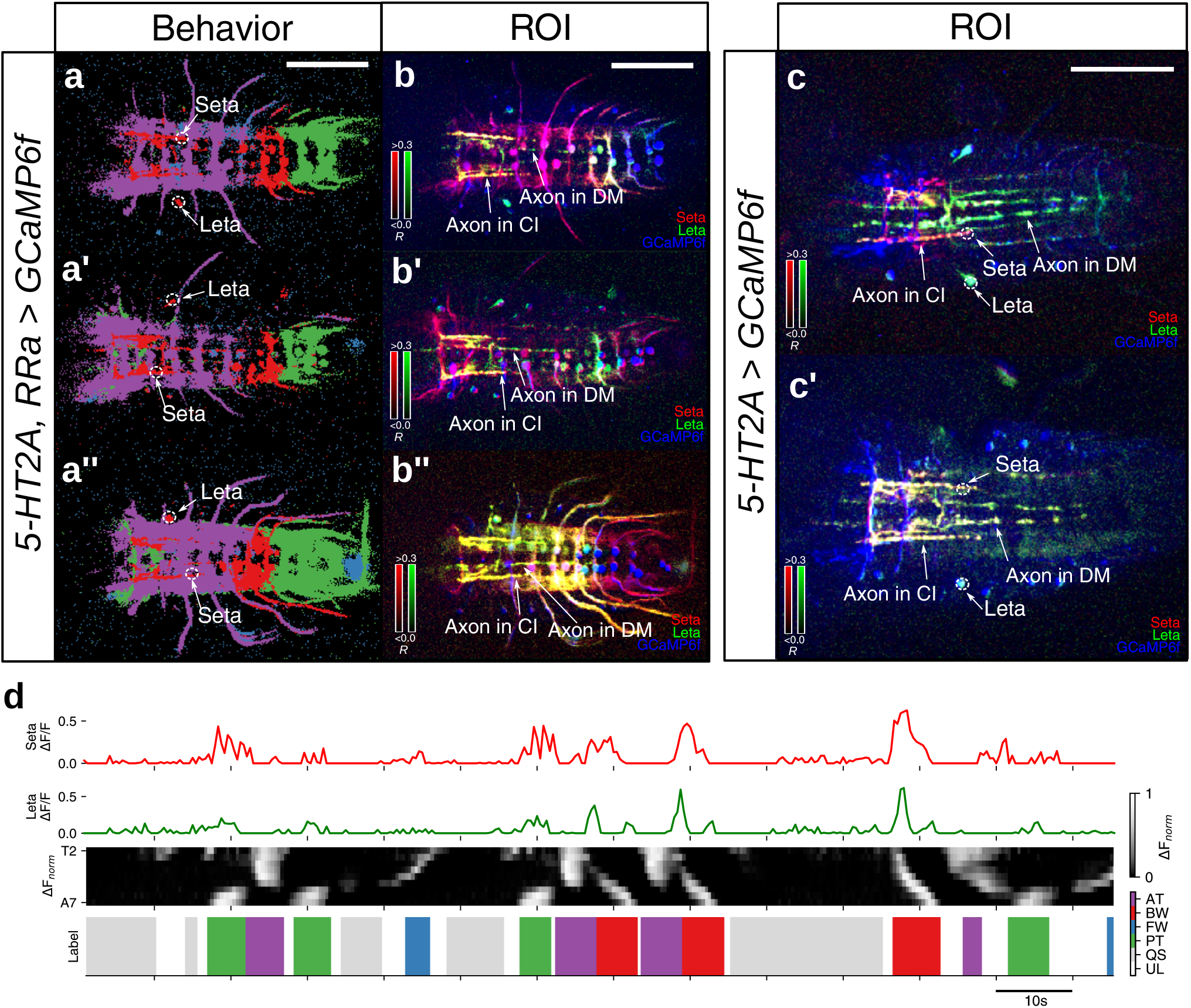
related to Fig. 4. (a-c) Dominant motor mapping (a-a”) and ROI correlation mapping (b-b”) as in Fig. 4c, and ROI correlation mapping (c, c’) as in Fig. 4f”, showing more samples. Note that a, b and c are the same as Fig. 4c’, C” and f”, respectively. (d) Activity of Seta and Leta neurons. Activity of single Seta (top, ∆F/F of Seta), and Leta (middle, ∆F/F of Leta) aligned with mapped motor patterns (bottom, ∆F*_norm_* of MNs and label). Color code of motor pattern as in Fig. 2. Scale bar 100 *µ*m.

**Figure S5.**
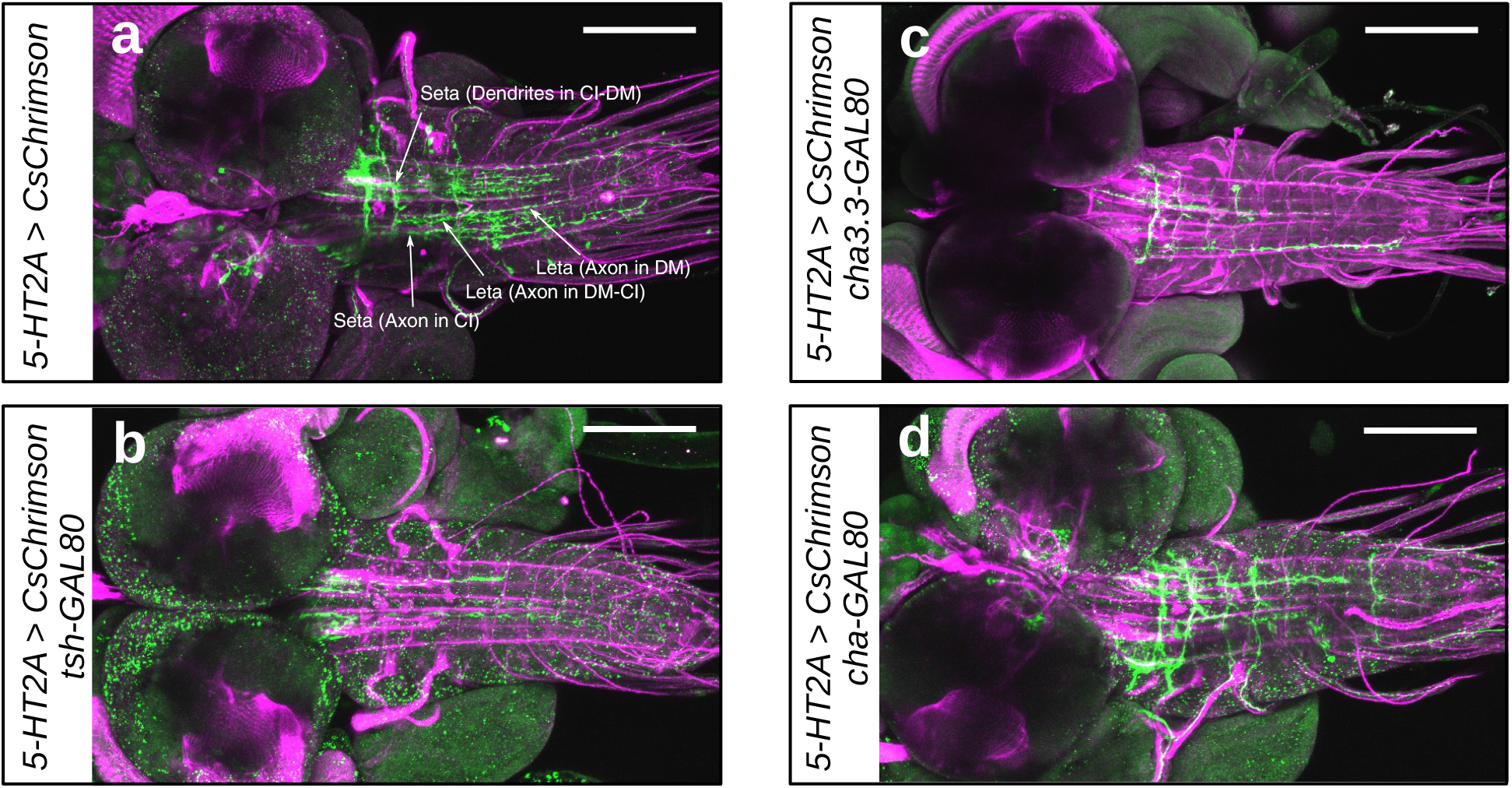
related to Fig. 6. Expression of CsChrimson driven by *5-HT2A-GAL4* (a) and its partial suppression by *tsh-GAL80* (b), *cha3.3-GAL80* (c) and *cha-GAL80* (d). Expression in Seta and Leta is absent in (b-d). Scale bar 100 *µ*m.

**Figure S6.**
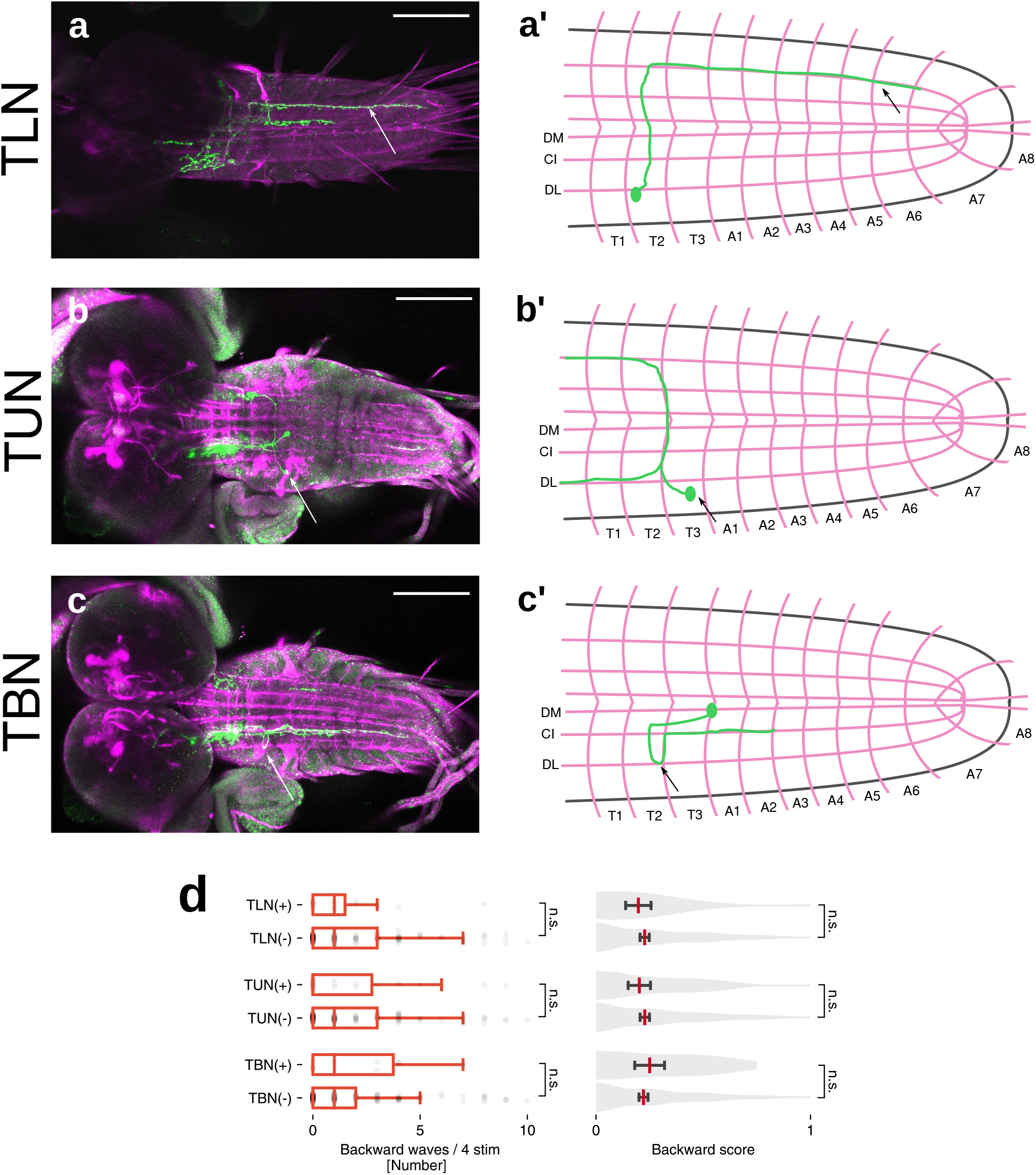
related to Fig. 6. Morphology of *5-HT2A-GAL4*-targeted neurons used as controls. (a-c) Immunostaining (green, a-c) and schematic diagram showing the morphology of TLN (a, a’), TUN (b, b’) and TBN (c, c’). Fas2 landmarks are also shown in a-c (magenta). (d) Quantification of the number of backward peristalsis (left, boxplot with scatter) and backward score (right, violinplot with mean and SEM) in larvae with or without CsChrimson expression in TLN, TUN and TBN. p=0.35 (waves), 0.36 (score) for TLN (n=19 (+), 161 (-)), p=0.44 (waves), 0.32 (score) for TUN (n=26 (+), 154 (-)), p=0.24 (waves), 0.29 (score) for TBN (n=14 (+), 166 (-)). P-value is derived from Mann–Whitney *U* test. Scale bar 100 *µ*m.

**Figure S7.**
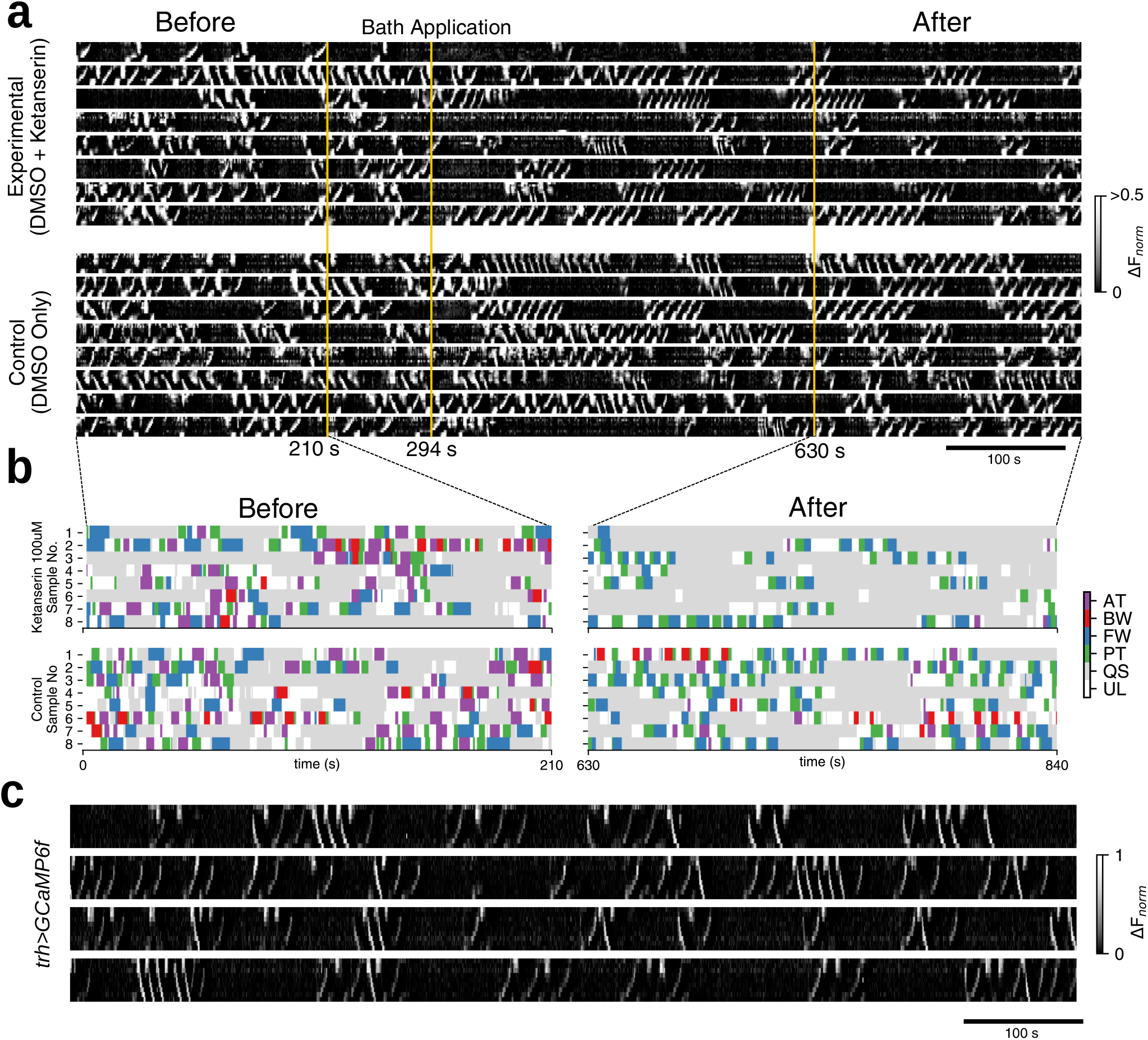
related to Fig. 7. (a-b) Segmental activity (T2-A7 from top to bottom in each row, ∆F*_norm_*) (a) and extracted motor pattern (b) before and after the application of Ketanserin obtained from 16 *5-HT2A,RRa>GCaMP6f* larvae. (c) Serotonergic neurons show stronger activity during BW and AT compared to during FW and PT. Segmental activity pattern (T2-A7 from top to bottom in each row, ∆F*_norm_*) from four *trh>GCaMP6f* larvae.

**Video S1.**
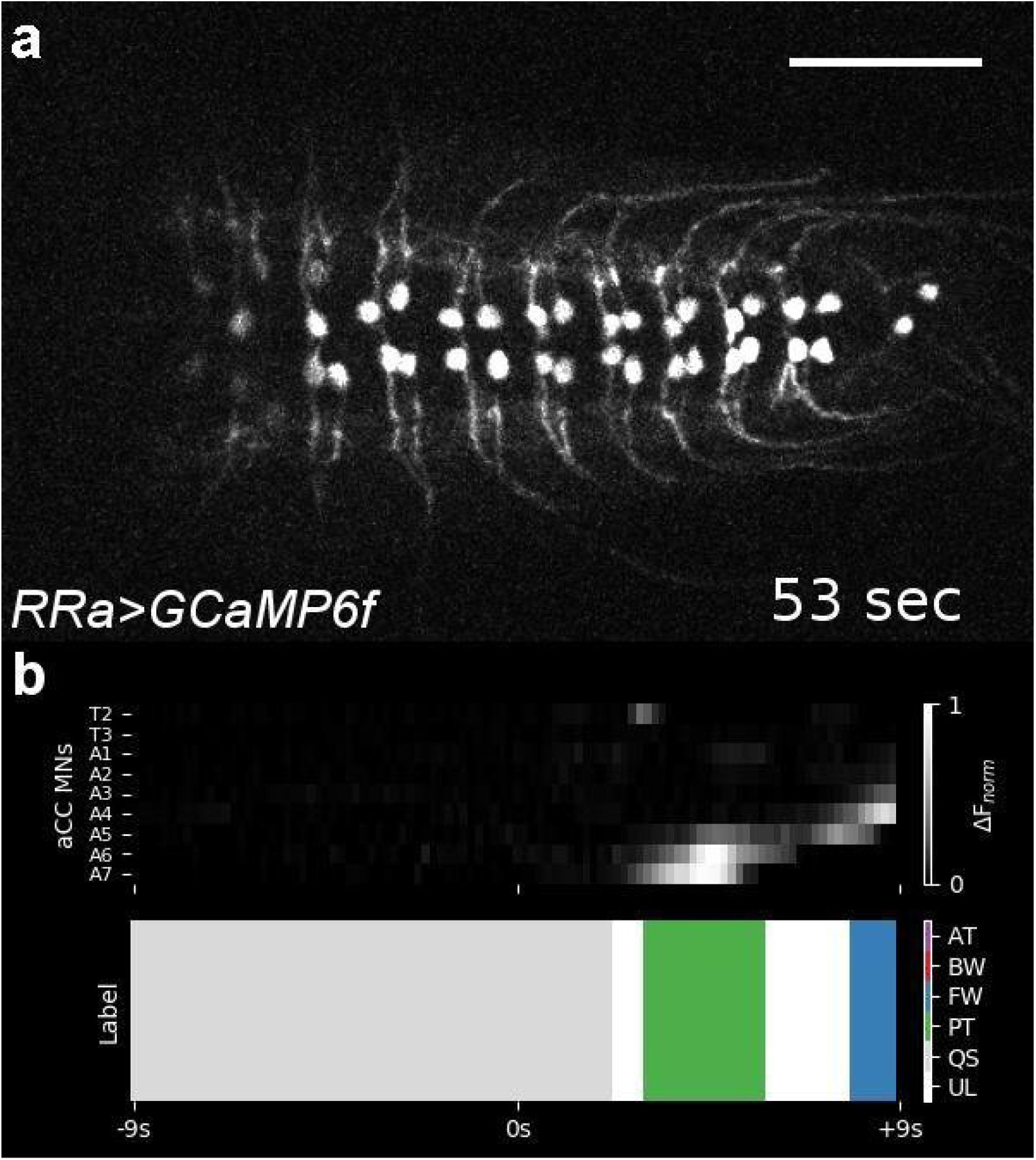
related to Fig 2. a’, a”’. (a) Calcium imaging movie of a sample of *RRa>GCaMP6f* with 5.4x speed. (b) pre-processed ∆F*_norm_* of 9-d timeseries displayed as a grayscale image (top) and label (bottom). Scale bar 100 *µ*m.

**Video S2.**
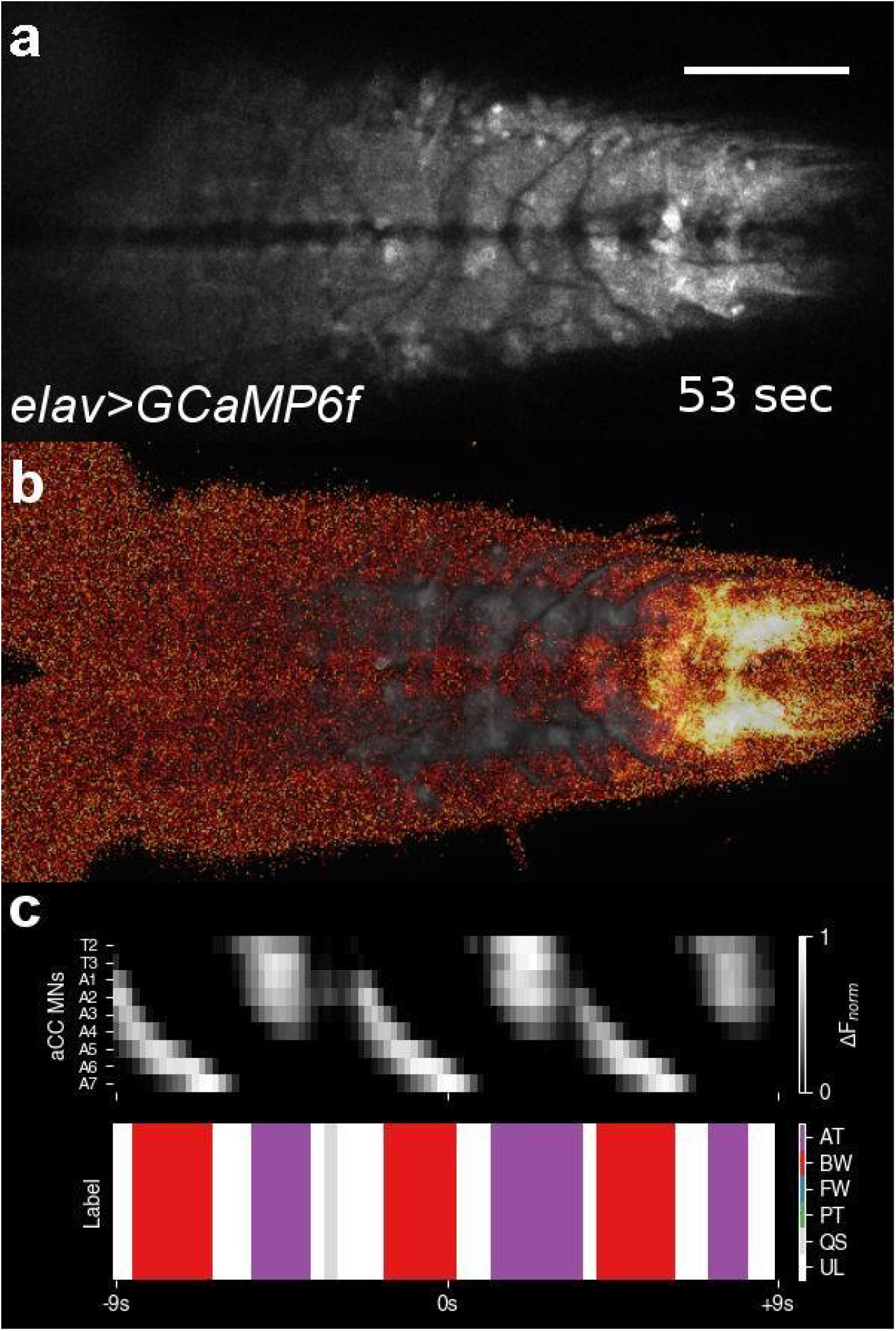
related to Fig. 3a. (a) Calcium imaging movie of a sample of *elav>GCaMP6f* with 5.4x speed. (b) Voxel-wise smoothed movie of (a). (c) pre-processed ∆F*_norm_* of 9-d timeseries displayed as a grayscale image (top) and label (bottom). Scale bar 100 *µ*m.

**Video S3.**
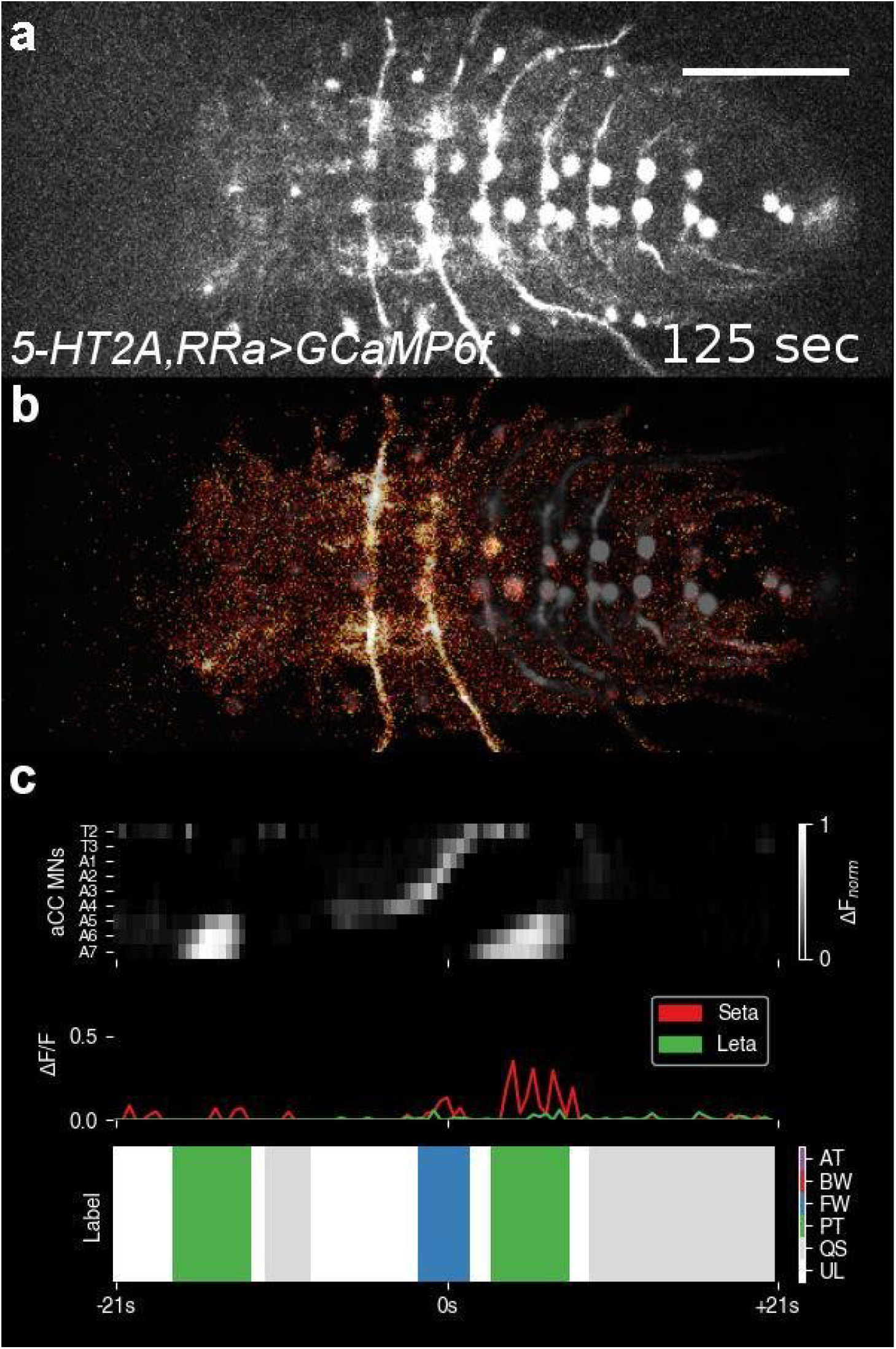
related to Fig. 4a. (a) Calcium imaging movie of a sample of *5-HT2A, RRa>GCaMP6f*with 12.6x speed. (b) Voxel-wise smoothed movie of (a). (c) pre-processed ∆F*_norm_* of 9-d timeseries displayed as a grayscale image (top), ∆F/F of Seta/Leta (middle) and label (bottom). Scale bar 100 *µ*m.

**Video S4.**
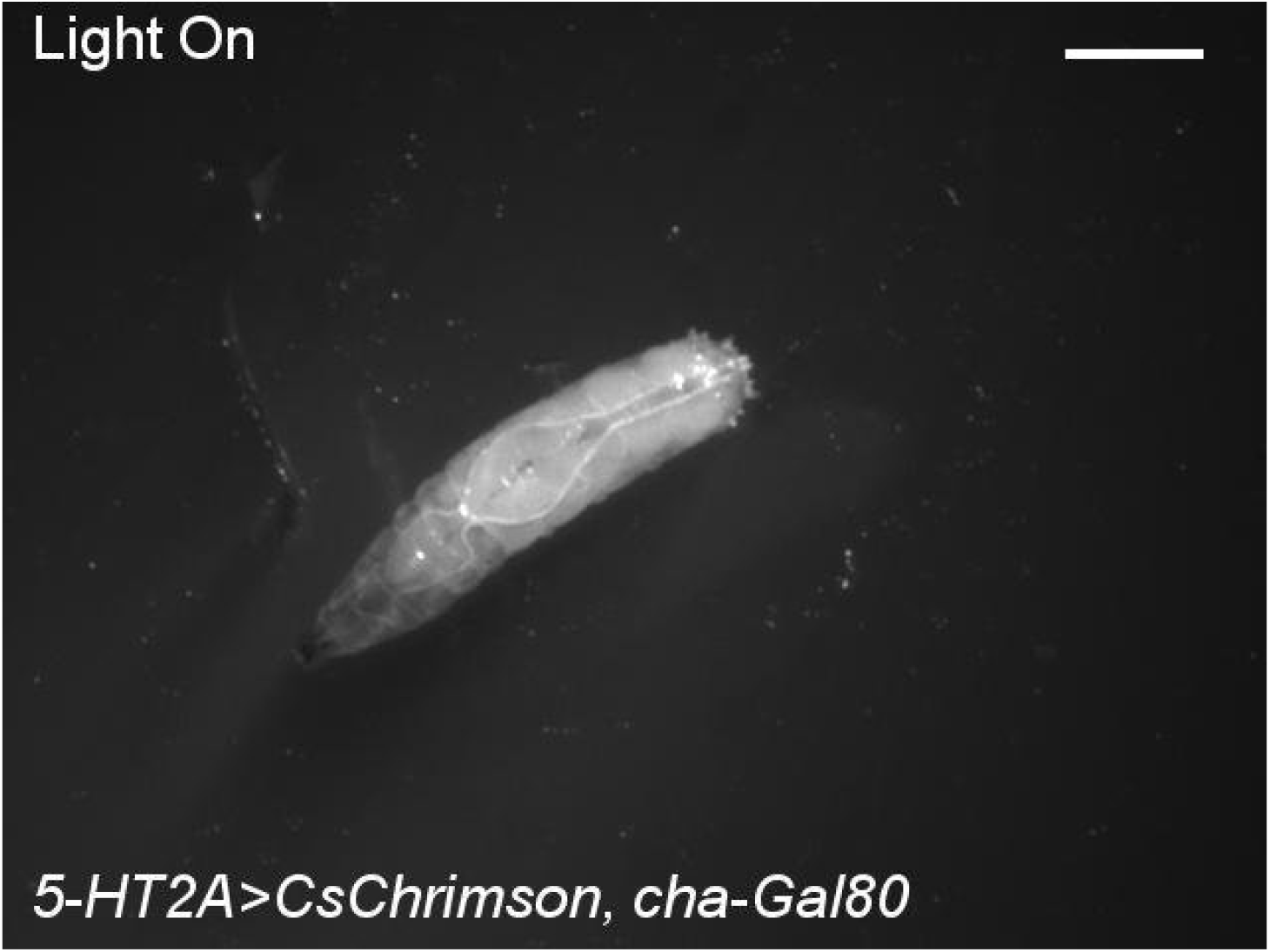
related to Fig. 6b. Optogenetic activation of *5-HT2A>CsChrimson* and *5-HT2A>CsChrimson*. 2x speed. Scale bar 1mm.

**Video S5.**
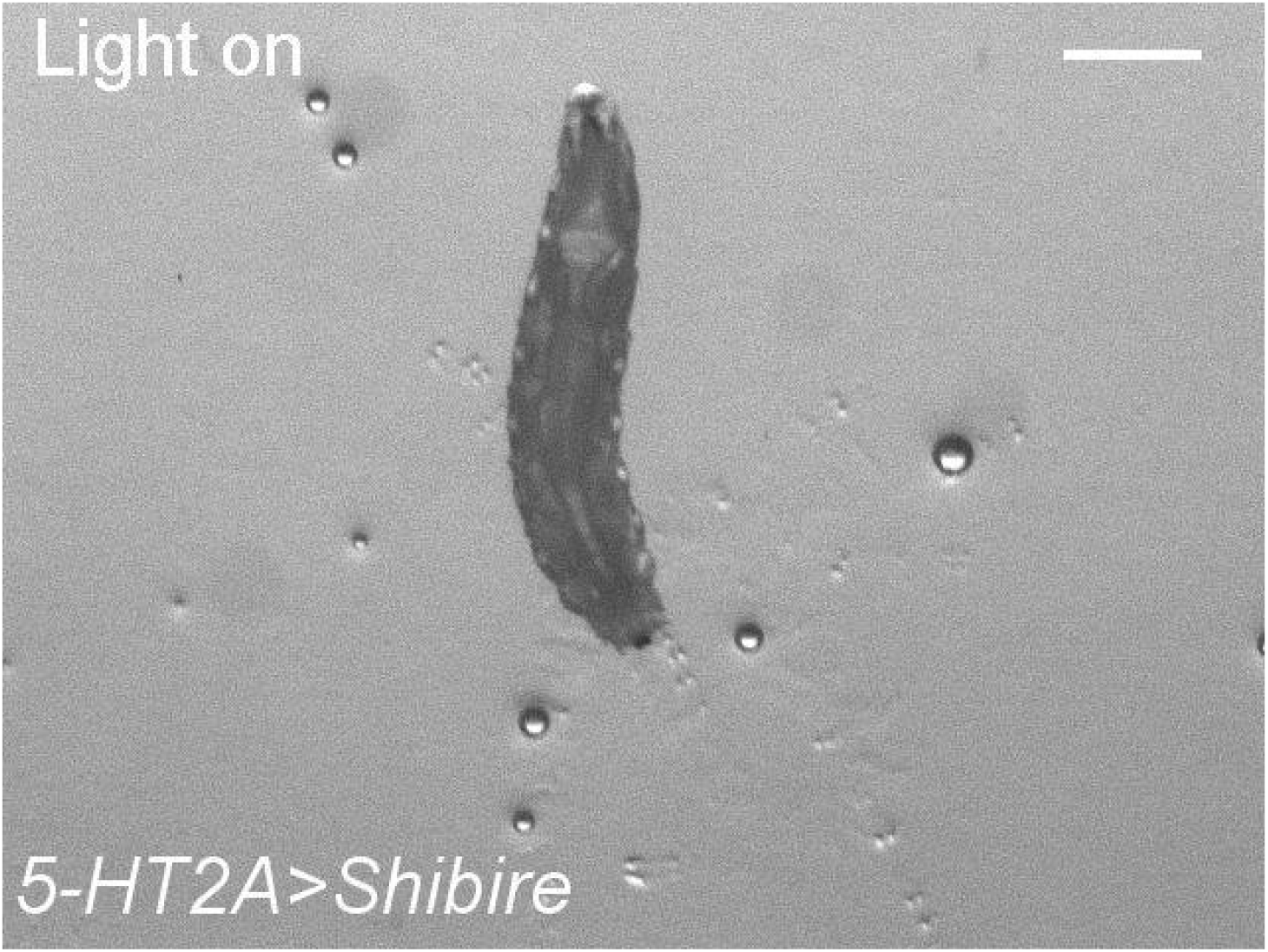
related to Fig. 7a. Blue light stimulation of *UAS-Shibire*, *5-HT2A>Shibire*, 2x speed. Scale bar 1mm.

**Video S6.**
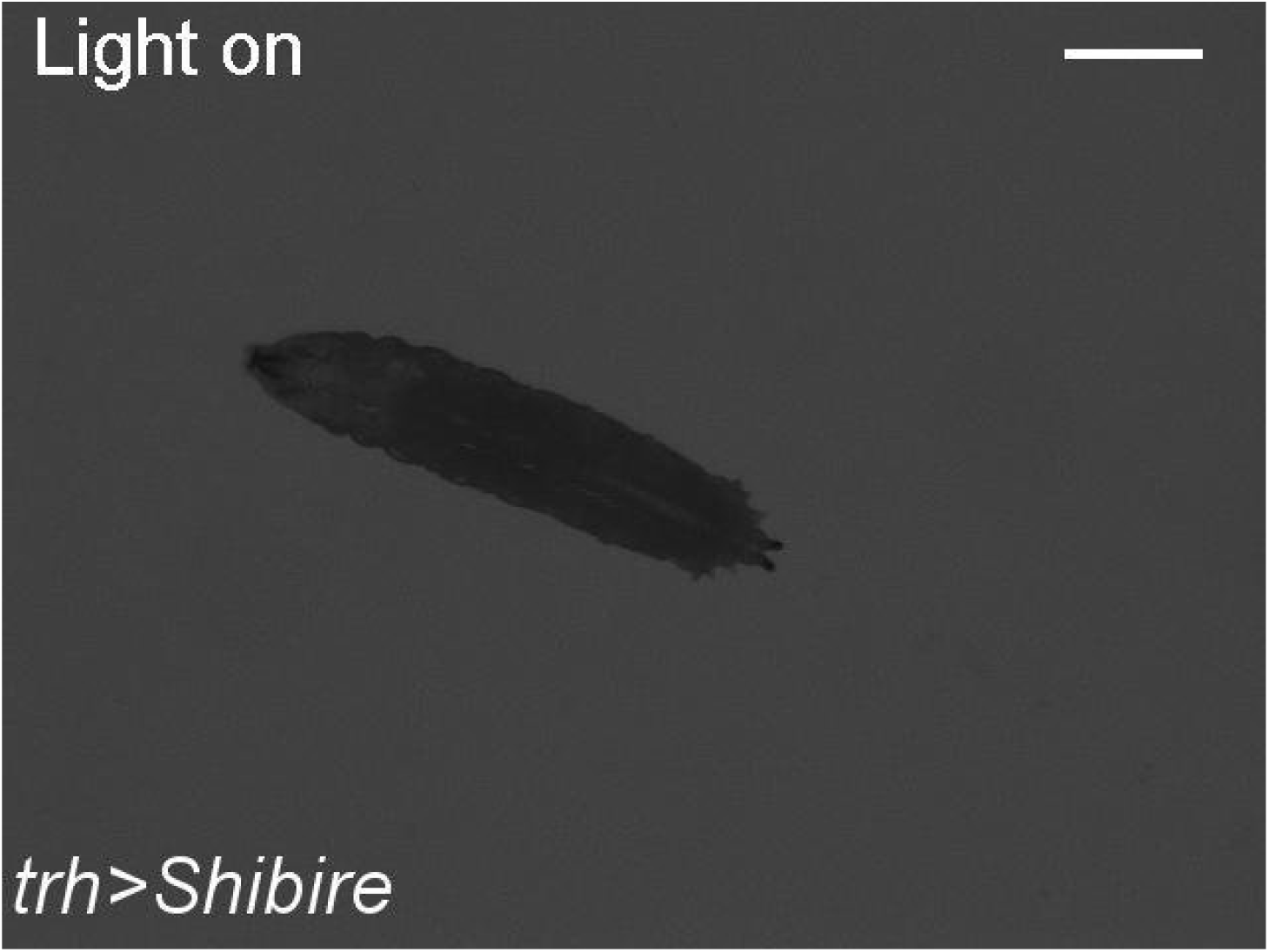
related to Fig. 7a. Blue light stimulation of *trh-GAL4* and *trh>Shibire*. 2x speed. Scale bar 1mm.

**Video S7.**
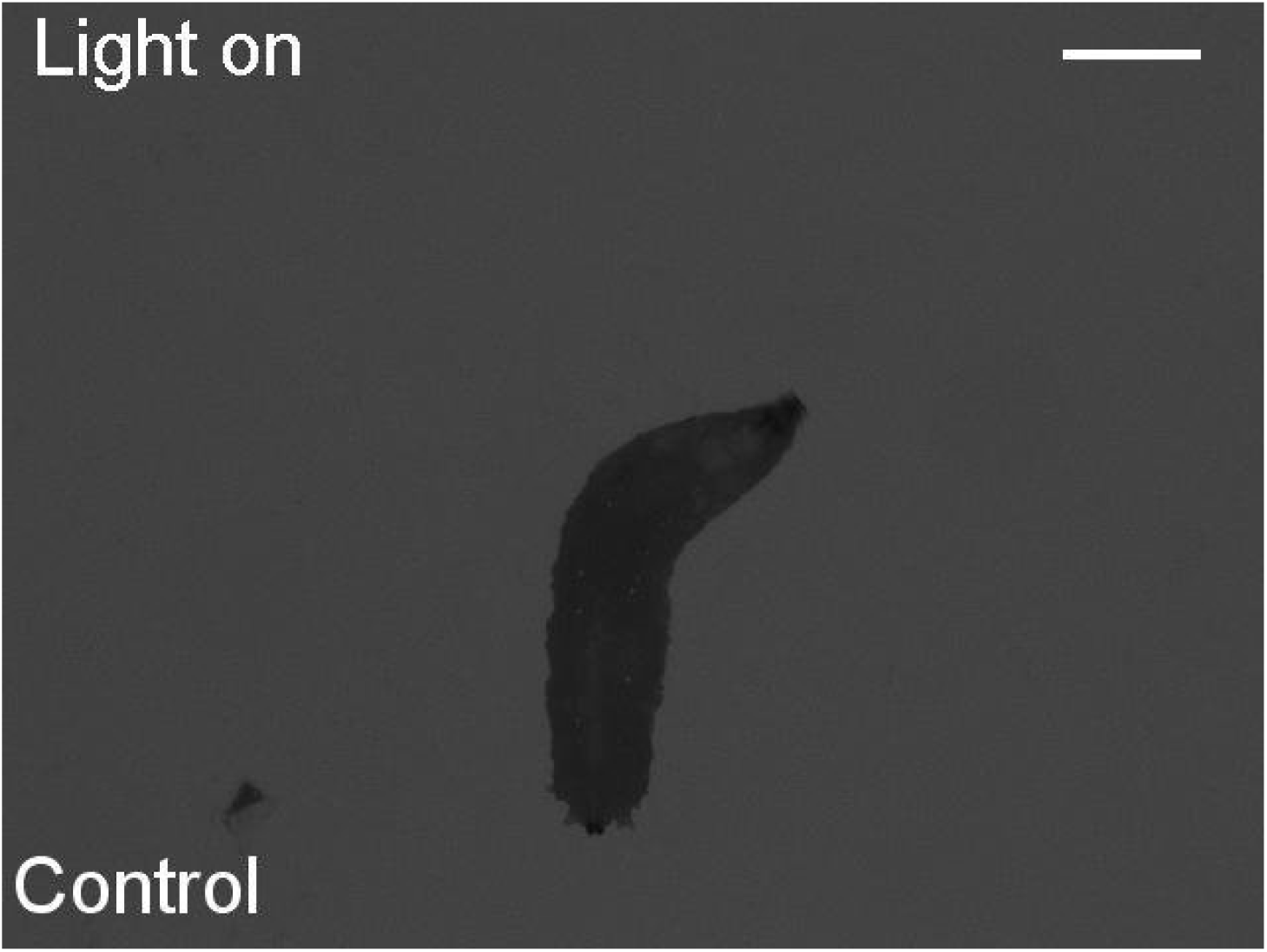
related to Fig. 7b. Blue light stimulation of *5-HT2A^PL^*. 2x speed. Scale bar 1mm.

**Video S8.**
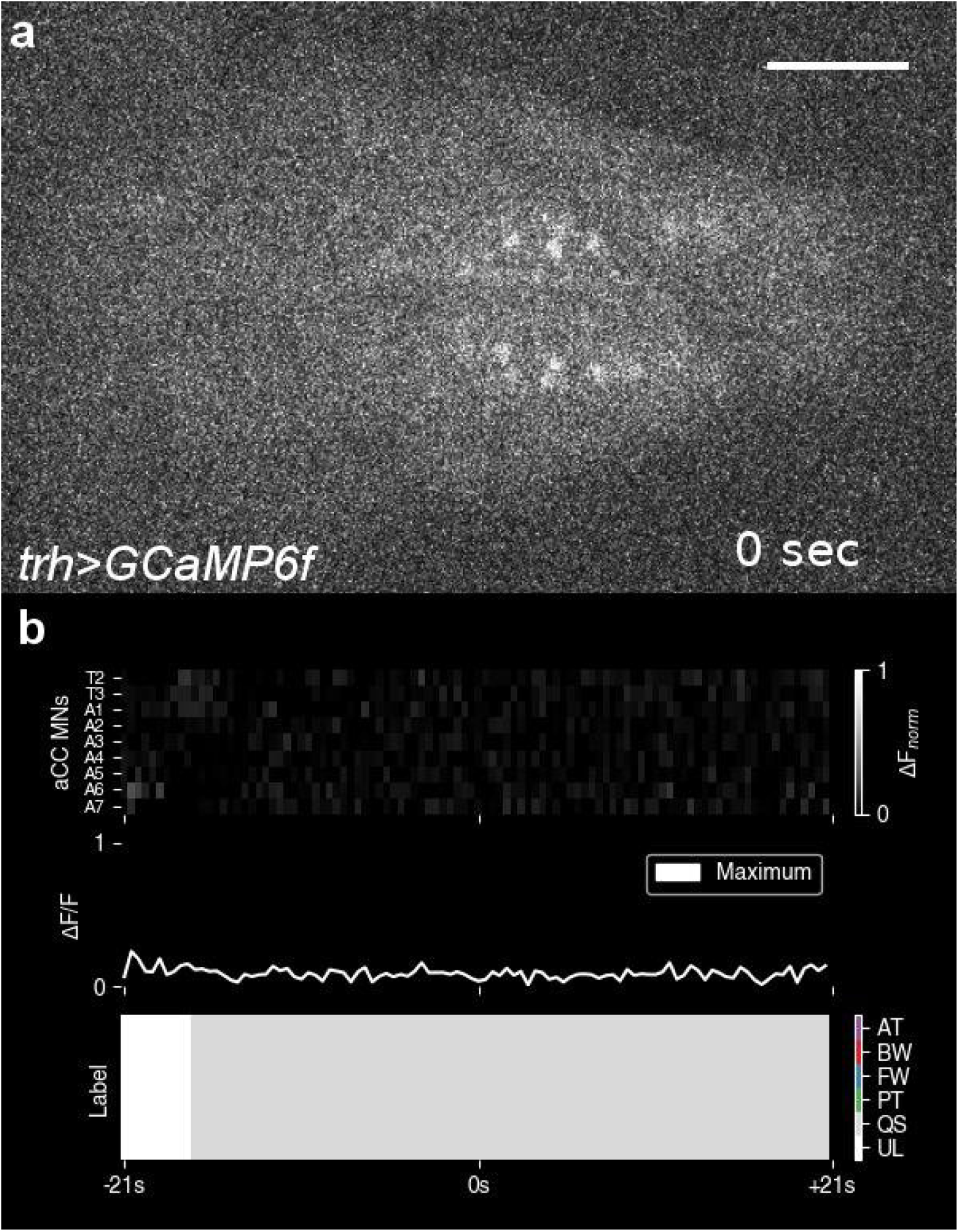
related to Fig. 4a. (a) Calcium imaging movie of a sample of *trh>GCaMP6f* with 12.6x speed. (c) pre-processed ∆F*_norm_* of 9-d timeseries displayed as a grayscale image (top), Maximum ∆F/F within ROIs (middle) and label (bottom). Scale bar 100 *µ*m.

